# Recombinant dimeric PDZ protein inhibitors for long-term relief of chronic pain by AAV therapeutics

**DOI:** 10.1101/2023.03.03.530962

**Authors:** Gith Noes-Holt, Kathrine L. Jensen, Mette Richner, Raquel Comaposada-Baro, Line Sivertsen, Sara E. Jager, Lucía Jiménez-Fernández, Rita C. Andersen, Jamila H. Lilja, Andreas H. Larsen, Marco B. K. Kowenicki, Sofie P. Boesgaard, Grace A. Houser, Nikolaj R. Christensen, Anke Tappe-Theodor, Christian B. Vægter, Rohini Kuner, Kenneth L. Madsen, Andreas T. Sørensen

## Abstract

The inadequate state of current pain treatments, the chronic nature of particularly neuropathic pain, and the high impact on quality of life render chronic pain conditions relevant for gene therapy. Here, we describe the development and application of self-assembling dimeric peptide inhibitors of the pain-associated scaffolding protein PICK1 (protein interacting with C-kinase 1) delivered by adeno-associated viral (AAV) vectors. In mice, these peptides prevented mechanical allodynia in inflammatory and neuropathic pain models and reversed neuropathic pain in advanced stages up to one year. Pain relief was obtained by targeting several relays along the somatosensory pain pathways unaccompanied by overt adverse side effects, while selective transduction of peripheral neurons was sufficient for providing full pain relief. We further confirmed PICK1 expression and peptide target engagement in mice and human donor tissue, and we conclude that AAV therapeutics, based on recombinant PICK1 inhibitors, represent a potential clinically meaningful strategy for persistent neuropathic pain conditions.

**One Sentence Summary:** Alleviating neuropathic pain by PICK1-directed gene therapy.

## INTRODUCTION

Exploiting gene therapies for refractory pain conditions holds promise to overcome several adversities of current pain medicine, including low efficacy, dose-limiting side effects, and opioid addiction (*1, 2*). Today’s drawbacks not only inflict on clinical practice and patients’ quality of life but also come with severe socioeconomic consequences, which the present opioid crisis unquestionably confirms (*3*). It is estimated that one in five adults in the US is affected by chronic pain (*4*), of which approximately half suffer from chronic neuropathic pain (*5*). This condition arises from a lesion or disease of the somatosensory nervous system (*6*), and no single treatment is currently able to prevent or cure neuropathic pain.

The most applied, and considered the safest, gene therapy vector for nervous system application is the adeno-associated viral (AAV) vector. It is characterized by its replication-deficient, low immunogenic, and long-lasting expression profile without the need for repeated dosing (*2, 7*). Its innate ability to deliver therapeutic gene expression locally and towards a defined cell population adds powerful dimensions to AAV tailoring for disease treatment. While available treatments for chronic neuropathic pain exclusively target receptors and transporters (*8*), we envision that targeting intracellular pathways underlying the development of chronic pain conditions might be more efficacious and circumvent systemic side effects.

Synaptic PDZ (PSD-95/DLG-1/ZO-1) domain scaffold proteins are vital for the integrity and modulation of neuronal communication and therefore clinically relevant pain targets. More specifically, PDZ proteins regulate discrete and dynamic functions of signaling events, such as protein transport, ion channel signaling, and other signal systems through protein-protein interactions (*9, 10*). Several synthetic peptide inhibitors that can block specific PDZ domain interactions have been developed to modulate synaptic efficacy by interrupting such protein-protein interaction (*11–15*). The peptides are characterized by their C-terminal residues, typically 5-13 amino acids of length and with the identical motif of a known interacting partner. Notably, peptides that target PSD-95 (postsynaptic density protein 95) disrupt the NMDA-type ionotropic glutamate receptor-dependent signaling and can attenuate animal pain-related behaviors (*16, 17*). We recently addressed another PDZ scaffold protein, PICK1 (protein interacting with C-Kinase 1), as a potent target for pain modulation (*14, 18*). This protein is highly conserved, expressed in neuronal tissue across species, and best known for its trafficking role in regulating AMPA-type ionotropic glutamate receptor surface expression at synapses (*19–21*). We constructed a synthetic bivalent peptide targeting PICK1 having two C-terminal HWLKV-motifs fused by a PEG-linker and conjugated to a Tat-peptide, rendering the peptide cell-permeable (*14*). Intrathecal (i.t.) administration of this PICK1 targeting peptide, TPD5, fully reversed mechanical allodynia for 3 hours in the spared-nerve injury (SNI) model of neuropathic pain. Surprisingly, efficacy was only evident for a bivalent and not an analogous, monovalent peptide inhibitor (*14*).

In the current study, we conceived an AAV-based strategy that allows recombinant peptide inhibitors to post-translationally assemble into a dimeric configuration to mimic the necessary bivalent structure of TPD5. Two C-terminal HWLKV-motifs are thereby freely positioned to interact with PICK1. We assessed the therapeutic efficacy of this approach in mouse pain models and harnessed the ability of AAVs for effective and selective expression of the peptide towards neurons. We further combined this strategy with LoxP recombination approaches and AAV capsid technology to determine the site of action. Our data revealed that various pain modalities in both inflammatory and neuropathic pain models were completely prevented or reversed with a duration of at least one year after treatment and without causing overt side effects. Targeting either the PNS or CNS, we observed that both such interventions were sufficient to completely reverse pain hypersensitivity. Furthermore, we verified both PICK1 expression and the therapeutic peptides’ target engagement in mice as well as human tissue from both dorsal root ganglion (DRG) and spinal cord. These findings collectively reinforce PICK1 as a promising target for AAV gene therapies tailored to address refractory neuropathic pain conditions.

## RESULTS

### High-affinity recombinant dimeric peptides targeting PICK1 prevent inflammation-induced mechanical allodynia

Inspired by our previous work on a synthetic bivalent peptide inhibitor of PICK1, TPD5 (*14, 18*), we conceived a molecular design enabling posttranslational self-assembly of a monomeric peptide, encoded by the AAV vector, into a dimer. This gene cassette, consisting of an N-terminal dimerization domain originating from yeast basic-region leucine zipper (bZIP) transcription factor (GCN4), a flexible glycine linker region (GGGGS), and a C-terminal PICK1 PDZ-binding motif (HWLKV), was created in two configurations. One variant with the native aspartic acid (D) and serine (S) in positions 7 and 14, respectively, and another variant with two prolines (P) at the same positions within GCN4 (**Fig. 1A**) predicted to render the homo-dimerization more dynamic (**Fig. S1**). Biochemical experiments with biotinylated peptides, confirmed that the self-assembly of the peptides indeed demonstrated PICK1 binding affinity (GCN4-HWLKV, K_i,App_ = 119 nM; GCN4(7P14P)-HWLKV, K_i,App_ = 548 nM) (**Fig. S2A**) and that GCN4-HWLKV, but not GCN4(7P14P)-HWLKV, drove PICK1 oligomerization (**Fig. S2B-E**), similar to the bivalent peptide, TPD5 (*14*). After confirmation of PICK1 binding, we went on to show peptide expression in primary neuronal cell cultures. To obtain selective and constitutive pan-neuronal expression, the peptide gene cassette was driven by the human synapsin (hSyn) promoter. AAV vectors were manufactured in-house, including the two therapeutic vectors: AAV8-Di-C5 (AAV8-hSyn-GCN4-HWLKV) and AAV8-Di(7P14P)-C5 (AAV8-hSyn-GCN4(7P14P)-HWLKV). Primary mouse cortical cultures were transduced with either of these vectors or two negative control vectors: AAV8-tdTomato (AAV8-hSyn-tdTomato) and AAV8-Di-GS (AAV8-hSyn-GCN4-GGGGS-GGGGS), the latter designed as a non-binding variant. Subsequently, the cultures were processed for GCN4 immunocytochemistry to validate transgene peptide expression. Peptide expression was confined to the cytoplasm and excluded from the nucleus (**Fig. 1B**). We next tested the vectors pain relieving effect in the Complete Freund’s Adjuvant (CFA) model of inflammatory pain. The level of mechanical allodynia was determined as the paw withdrawal threshold to stimulation by von Frey filaments and was measured repeatedly before and after a single i.t. injection at the L5-L6 lumbar level in wild-type (WT) male mice. After approx. 3 weeks to allow for viral expression of the peptide, the mice were injected intraplantar with CFA to induce inflammatory pain in the left hind paw. Both the AAV8-Di-C5 and AAV8-Di(7P14P)-C5 vectors prevented the development of allodynia, whereas AAV-tdTomato injected control mice developed significant allodynia as expected in the days following intraplantar CFA injection (**Fig. 1C**). In addition, the pain threshold remained unaltered in the non-inflamed (contralateral) paw, regardless of treatments (**Fig. 1C**), suggesting an analgesic effect of the two active recombinant peptides without numbing (anesthetizing) mechanical sensitivity. After 11 days, the paw withdrawal threshold for AAV-tdTomato returned to previous baseline levels, highlighting the transient nature of this model.

**Fig. 1.**
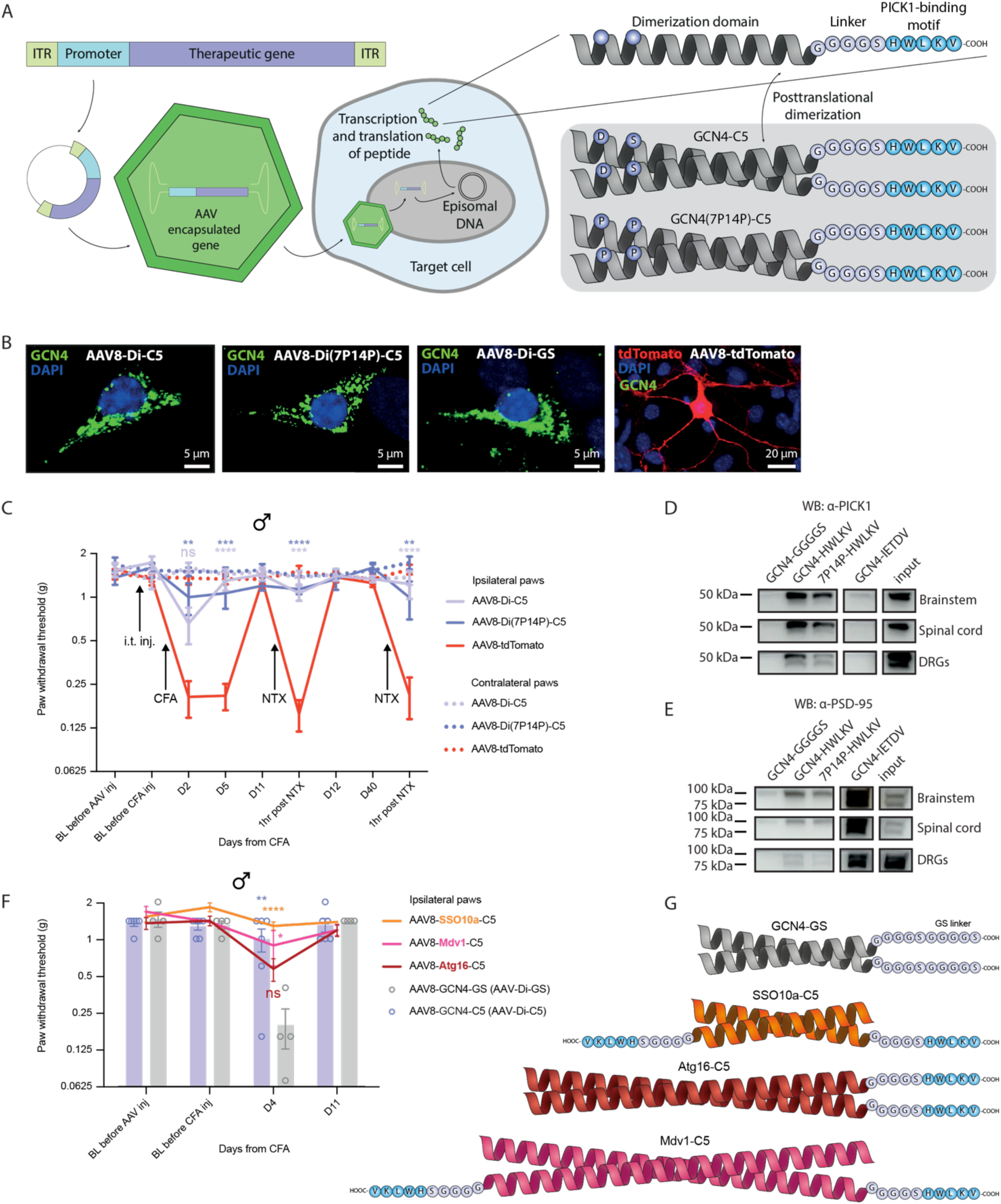
AAV-encoded dimeric peptides designed for targeting PICK1 fully abolish CFA-induced mechanical allodynia. **A)** Conceptual design of AAV-encoded dimeric peptide inhibitors with HWLKV motifs for targeting PICK1. Two different amino acid variants within the GCN4 helical structure are shown. **B)** ICC performed on cortical neuronal cultures transduced with different AAV variants, showing GCN4 immunoreactivity (green), DAPI nuclei staining (blue) and tdTomato autofluorescence (red) (for AAV-tdTomato only). **C)** Paw withdrawal threshold measured by von Frey before and after i.t. injection of AAVs, and again after CFA and NTX administration, as indicated. Ipsilateral paw withdrawal threshold of AAV8-tdTomato (control group) was compared to AAV8-Di-C5 and AAV8-Di(7P14P)-C5 treatment conditions, including the two EGFP containing vectors shown in Suppl. Figure 3A. \*\**P* < 0.01, \*\*\**P* < 0.001, \*\*\*\**P* < 0.0001. Two-way repeated measures ANOVA, F(32, 208) = 2.198; Dunnett’s multiple comparison; *N* = 6-8 per group. **D)** Western blot of PICK1 following pull-down from the brain stem, spinal cord, and DRG tissue homogenate spiked with biotinylated peptides. GCN4-HWLKV and GCN4(7P14P)-HWLKV peptides are designed to target PICK1, whereas GCN4-GS is a negative control, and GCN4-IETDV has a binding motif for targeting PSD-95. *N* = 2 mice (pooled). **E)** Western blot of PSD-95 in a similar setup as in panel D. **F)** Test of different AAV-encoded peptides in the CFA model measuring paw withdrawal threshold before and after i.t. injection and after CFA administration, where AAV8-Di-C5, AAV8-SSO10a-C5, AAV8-Mdv1-C5, AAV8-Atg16-C5, and AAV8-Di-GS (non-binding variant) treatments were compared. \**P* < 0.05, \*\**P* < 0.01, \*\*\*\**P* < 0.0001. Two-way repeated measures ANOVA, F(12, 57) = 2.449; Tukey’s multiple comparison; *N* = 4-6 per group. **G)** Illustration of AAV-encoded recombinant peptides variants, including the non-binding GCN4-GS variant and the three PICK1-binding variants (SSO10a, Atg16, Mdv1) having alternative dimerization domains of different lengths and orientations. Abbreviations: BL, baseline; i.t., intrathecal; NTX, naltrexone. All data are shown as mean with s.e.m.

Subsequently, the opioid receptor antagonist naltrexone (NTX) was administered once on day 11 and again on day 40 to reinstate allodynia (*22*). AAV8-Di-C5 and AAV8-Di(7P14P)-C5, but not AAV8-tdTomato treatment, prevented mechanical allodynia demonstrating an extended duration of action (**Fig. 1C**). The chronic nature of the AAV intervention renders it difficult to discern whether the treatment simply relieves the NTX-induced hyperalgesia or rather prevents the underlying plasticity of the opioid system so that NTX no longer reinstates hyperalgesia. AAVs with various EGFP tagging designs in combination with the therapeutic sequences were also efficacious in this CFA experiment (**Fig. S3A**). Next, we tested the effect of AAV8-Di-C5 and AAV8-tdTomato on heat-induced hyperalgesia. CFA injection induced a significant reduction in threshold temperature in AAV8-tdTomato injected animals, while this effect was smaller and non-significant for AAV8-Di-C5 injected mice. However, the threshold temperature for AAV8-Di-C5 mice was not significantly different from AAV8-tdTomato mice (**Fig. S3B**), suggesting a partial relief of heat hyperalgesia in addition to the full relief of mechanical allodynia.

To investigate target engagement and specificity, we performed pull-down experiments in mouse tissue using synthetic biotinylated GCN4-peptides, either containing the HWLKV motif designed for targeting PICK1, or the IETDV motif previously used to target PSD-95 (*13*). Western blots of peptide-spiked tissue lysates from the brain stem, spinal cord, and DRGs demonstrated robust target engagement and clear selectivity towards PICK1 and PSD-95, respectively (**Fig. 1D-E**). Finally, we asked whether the pain-relieving effects of the high-affinity PICK1 inhibitors were dependent on the GCN4 dimerization domain or the PDZ-binding sequence, HWLKV. First, we exchanged GCN4 with alternative dimerization domains; Atg16-C5, Mdv1-C5, and SSO10a-C5 having either longer lengths or antiparallel orientations (**Fig. 1F-G**). Atg16-C5 and Mdv1-C5 peptides were too long for synthetic production, but biochemical experiments with the SSO10a-C5 peptide confirmed that antiparallel self-assembly mediated by the SSO10a helix conferred affinity for PICK1 binding and drove PICK1 tetramerization similar to GCN4-C5 (**Fig. S4A-E**). Secondly, we substituted the PDZ-binding sequence, HWLKV, with a second copy of the linker sequence, GGGGS, to yield GCN4-GS. Biochemical experiments validated the intact helical and dimeric assembly of GCN4-GS, while PICK1 binding and tetramerization were abolished (**Fig. S4A-D, F**). Upon i.t. administration of AAV8-GCN4-GS, mice developed mechanical hyperalgesia in the CFA model similar to the negative control, AAV8-tdTomato (**Fig. 1F**). In contrast, injection of AAV8-Di-C5 again significantly alleviated mechanical allodynia, demonstrating the critical role of the PDZ-binding motif (**Fig. 1F-G**). Injection of AAV8-Atg16-C5, AAV8-Mdv1-C5, or AAV8-SSO10a-C5 relieved mechanical allodynia to varying extents, with AAV8-Atg16-C5 not being significantly different from AAV8-GCN4-GS (**Fig. 1F-G**). This demonstrates that the observed treatment effects were not dependent on the GCN4 motif itself. In contrast, the varying effect of the substitution for GCN4 might reflect differences in dimerization strength, steric hindrance, expression level, etc. To this end, we concluded that different high-affinity recombinant dimeric peptides, targeting PICK1 and delivered by AAV, can prevent mechanical allodynia in the CFA model, even during NTX-induced hyperalgesia reinstatement suggesting a non-opioid analgesic profile on nociceptive pain. We decided, therefore, to further characterize the expression and efficacy profiles of AAV8-Di-C5 and AAV8-Di(7P14P)-C5 vectors.

### Targeting neurons below the cervical spinal level can reverse mechanical allodynia

To determine which neurons contributed to the pain-relieving effect following i.t. AAV administration, we took advantage of the transgenic mouse line, Hoxb8-Cre. This mouse expresses Cre recombinase under the control of the *Hoxb8* gene and is confined to neurons caudal to the cervical level 3 of the spinal cord, including the DRGs (*23*). Cre-dependent AAVs were designed to achieve confined peptide expression by exploiting a Cre-ON (AAV8-DIO[EGFP-P2A-Di-C5]ON) and Cre-OFF (AAV8-DIO[EGFP-P2A-Di-C5]OFF) approach (**Fig. S5A-B**). We again used the CFA model to first confirm the robustness of the AAVs, now in mice of mixed genders. In WT mice, the OFF-variant, designed to express the Di-C5 peptide constitutively in non-Cre expressing neurons, fully prevented the development of mechanical allodynia (**Fig. S5C-D**) in agreement with our previous result (**Fig. 1C**). As expected, the ON-variant was in contrast without effect in WT animals (**Fig. S5C-D**). However, when tested in Hoxb8-Cre mice, the Cre-ON variant (i.e. peptide expression caudally to C3) fully abolished mechanical allodynia. The Cre-OFF variant (i.e. peptide expression rostrally to C3), on the other hand, displayed a trending, but non-significant treatment effect (**Fig. S5C-D**). From this, we conclude that transgene expression caudally to cervical level 3 is adequate to mitigate mechanical allodynia.

To obtain insight into the transduction profile following i.t. delivery of the recombinant AAV, we next used the Cre-reporter mouse line, Ai14, that expresses robust tdTomato fluorescence following Cre-mediated recombination (**Fig. S6A)**. For this purpose, we employed an AAV similar to AAV8-EGFP-P2A-Di-C5 (**Fig. S3A),** but now additionally encoding iCre fused to EGFP (AAV8-iCre:EGFP-P2A-Di-C5) to induce tdTomato expression in transduced neurons. Tissues were examined 40-44 days after AAV delivery, and both cryostat-prepared sections and cleared whole-mount spinal cords were imaged. For the latter, we extended the iDISCO tissue-clearing technique with decalcification, rendering both soft and hard tissues transparent and suitable for visualization using light-sheet microscopy. Clear tdTomato signal was detectable within neuronal cell bodies of DRGs at sacral, lumbar and thoracic segments (**Fig. 2A-C, Fig. S6B**) with visible ascending fibers confined within the spinal cord dorsal column (**Fig. 2D, Fig. S6B-C**) consistent with robust transduction of mechanosensitive Aβ fibers. This labeling pattern was accompanied by thin fibers extending into the spinal cord dorsal horn laminae (**Fig. 2D**), although is unclear whether these fibers are of nociceptive origin. The tdTomato fluorescence along the spinal cord terminated at the dorsal column nuclei in the closed medulla oblongata (cuneate and gracile nuclei) and did not extend further rostrally from the pons (**Fig. 2E**). However, sparse labeling of individual neuronal cell bodies was observed in the brain, e.g., in the cerebellum and olfactory structures (**Fig. S6C**). Little if any expression was observed in cell bodies within the spinal cord at the lumbar level (**Fig. 2A-B, D**). This expression pattern suggests that AAV8, after i.t. injection, predominantly transduces DRGs having fibers extending into local spinal circuits but with noticeable projections extending to supraspinal sites.

**Fig. 2.**
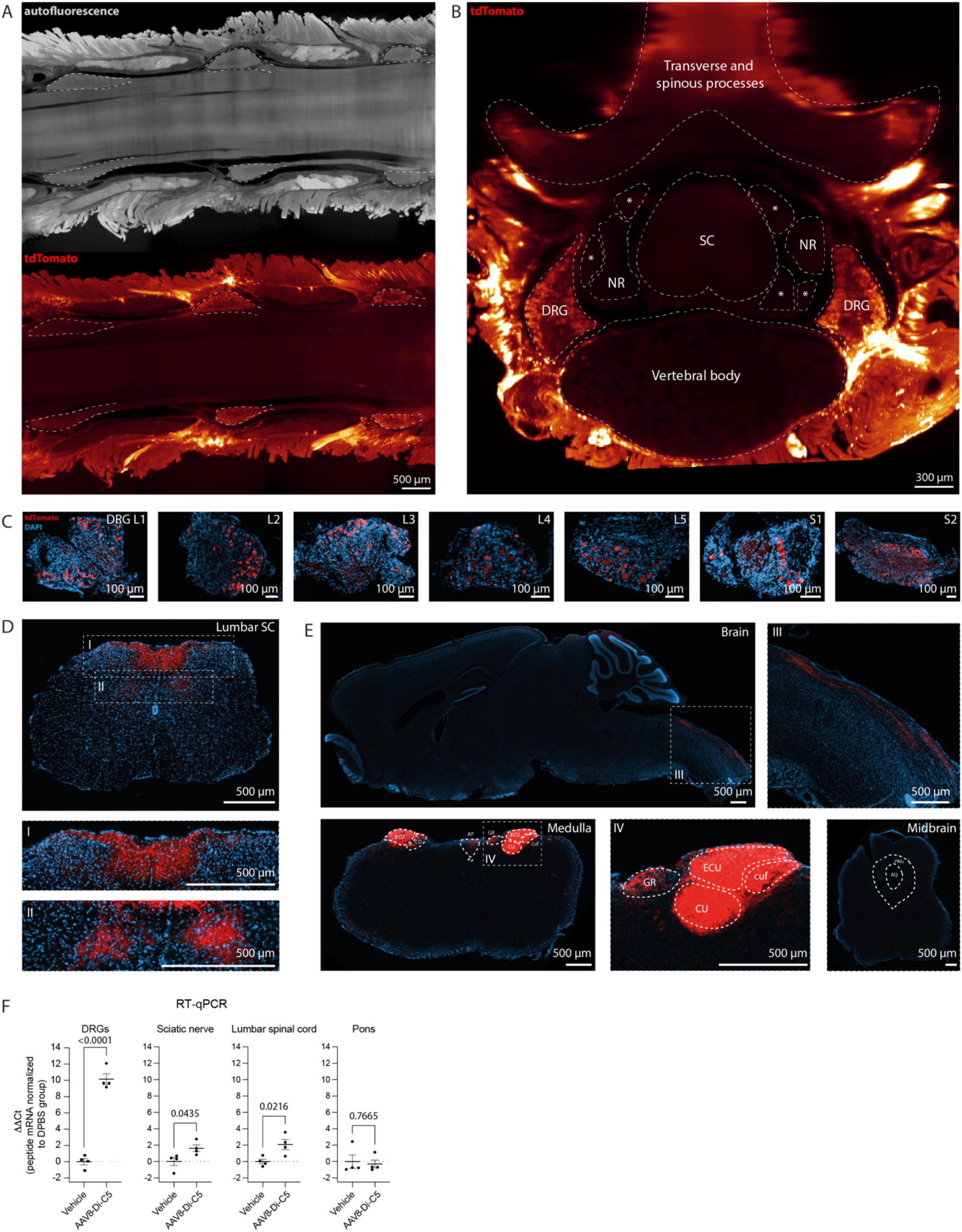
Peptide expression following intrathecal AAV8 injection is primarily within dorsal root ganglion neurons. A) Longitudinal view using whole-mount imaging showing the lumbosacral spinal cord region labeled by tdTomato following i.t. injection of AAV8-iCre:EGFP-P2A-Di-C5 vector into Ai14 reporter mice. Top image, shown by the autofluorescence signal, outlines the spinal cord and pairs of DRGs (dotted lines). Bottom image, in the same plan as above, shows tdTomato labeled neurons within DRGs. B) Transverse view of the spinal cord displayed in panel A. DRGs on both sides are labelled, with no neuronal somas detectable within the spinal cord. Non-neuronal tissue surrounding the vertebrae column is also labeled by tdTomato. Abbreviation: NR, nerve root (also indicated by asterisk). *N* = 8 mice for panel A-B. C) Cryostat sections showing tdTomato-labeled neurons within lumbar (L1-L5) and sacral (S1-S2) DRGs. D) Transverse section of the spinal cord at the lumbar level with notable tdTomato labeled projections within the dorsal column spinal cord comprising ascending fibers from DRG neurons (Insert I). Within the grey matter of the dorsal horn, fibers, but no neuronal somas, were visible (insert II, gain adjusted). E) Sagittal brain section with tdTomato-positive fibers within the brain stem (Insert III) terminating within cuneate and gracile nuclei (Insert IV) and not extending rostrally into periaqueductal gray (PAG) and beyond. The bottom three panels are gain adjusted. Abbreviations: AP, area postrema; AQ, cerebral aqueduct; ECU, external cuneate nucleus; CU, cuneate nucleus; cuf, cuneate fascicle; GR, gracile nucleus. *N* = 9 mice for panel C-D. F) Peptide mRNA expression levels in DRGs from L3-L5 (both sides), sciatic nerve, lumbar spinal cord, and pons evaluated by RT-qPCR. Naïve WT mice received i.t. injection of either AAV8-Di-C5 or vehicle (DPBS). Unpaired t-test, DRGs: t(6) = 13.20; Sciatic nerve: t(6) = 2.549; Lumbar spinal cord: t(6) = 3.084; Pons: t(6) = 0.3108. *N* = 4 mice per group. Data are shown as mean with s.e.m.

The Ai14 mice of mixed gender also did not develop mechanical allodynia in the CFA model following i.t. delivered AAV8-Di-C5, suggesting that these tdTomato-expressing neurons are sufficient for pain relief (**Fig. S6D)**. Further examination revealed that transduced tdTomato-positive DRG neurons, which were stained for IB4, CGRP, or NF200 markers, comprised a fraction of all DRGs (**Fig. S6E-F**) and were significantly shifted towards larger DRG neurons (**Fig. S6F**). While smaller DRG neurons consisted mainly of IB4+ or CGRP+ neurons, the larger DRG neurons mainly comprised CGRP+ or NF200+ neurons (**Fig. S6G**). In a separate cohort of WT mice, we saw that mRNA level of the peptide was notably high in DRGs of AAV8-Di-C5 injected mice (**Fig. 2F, Table S1**), in agreement with the tdTomato signal seen in Ai14 mice (**Fig. 2A-D)**. In the sciatic nerve and lumbar spinal cord, mRNA peptide levels were also detectable, but not equally prominent. In all other tissues examined, except for the liver, peptide mRNA was not identified (**Fig. 2F, Table S1)**. We concluded that i.t. delivery of our AAV therapeutics give rise to significant DRG expression, which is sufficient to relieve mechanical allodynia.

### Complete prevention and reversal of chronic neuropathic pain by AAV-encoded PICK1 inhibitors

Having identified the DRGs as site of action sufficient for pain relief, we next explored the efficacy of the AAVs in the SNI model of neuropathic pain in three different treatment paradigms. In the first treatment paradigm, we evaluated the recombinant peptides in preventing the occurrence of mechanical allodynia. WT mice were i.t. injected 14 days before SNI surgery, and mechanical allodynia was again determined by paw withdrawal threshold using von Frey filaments. Following the SNI procedure, AAV8-tdTomato, as well as vehicle (DPBS), treated male mice showed robust and comparable levels of mechanical allodynia until at least 28 days after SNI, whereas AAV8-Di-C5 and AAV8-Di(7P14P)-C5 completely prevented allodynia as compared to AAV8-tdTomato and vehicle (**Fig. 3A**). In a similar setup, AAV8-Di(7P14P)-C5 was equally effective in female mice (**Fig. 3B**). These findings prompted us to further assess the treatment efficacy and robustness in the SNI model. In a second treatment paradigm, AAVs were delivered five weeks following SNI injury, at a stage when initial inflammation has ceased. When comparing AAV8-Di(7P14P)-C5 and AAV8-tdTomato treated mice, mechanical allodynia was significantly attenuated from week 9 after SNI and until the end of the experiment at day 173 when comparing AAV8-Di(7P14P)-C5 and AAV-tdTomato-treated male mice (**Fig. 3C**). Finally, in the last treatment paradigm, we assessed the long-term efficacy of the intervention, where male SNI mice were repeatedly monitored for an entire year. With i.t. injections performed on day 2 after injury, AAV8-Di(7P14P)-C5 significantly reversed mechanical allodynia for 365 days, except at day 322, as compared to AAV8-tdTomato (**Fig. 3D**). Based on these findings, we concluded that the recombinant peptides can prevent the development and reverse an already established mechanical allodynia in both acute and chronic stages of neuropathic pain and maintain efficacy for at least 1 year in mice.

**Fig. 3.**
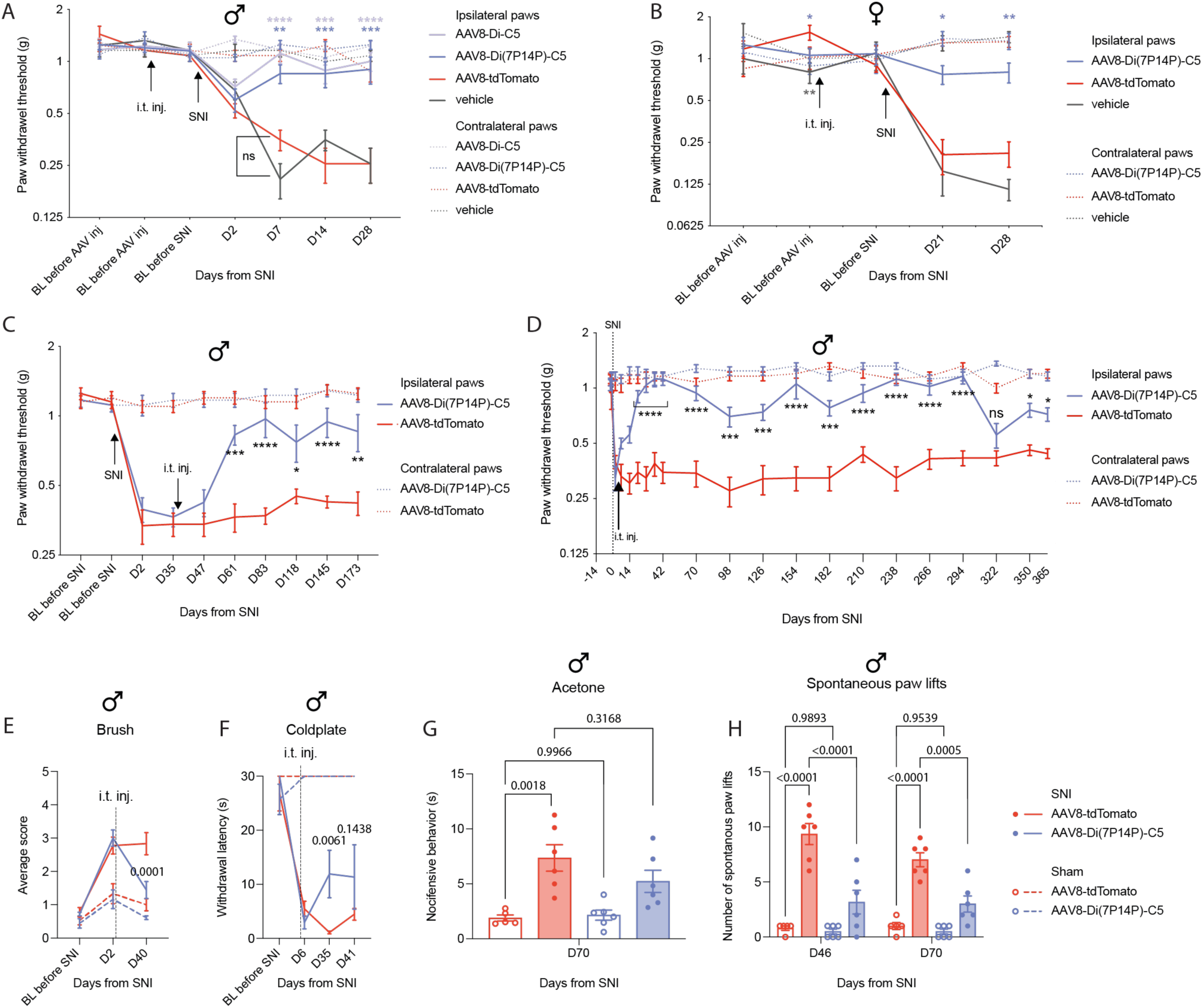
AAV-encoded dimeric peptide targeting PICK1 revokes SNI-induced allodynia for at least one year. **A)** Paw withdrawal threshold in WT male mice before and after i.t. injection and after SNI in a preventative treatment paradigm, where AAVs or vehicle (DPBS) are administered before SNI surgery. \*\**P* < 0.01, \*\*\**P* < 0.001, \*\*\*\**P* < 0.0001. Two-way repeated measures ANOVA, F(18, 126) = 5.474; Tukey’s multiple comparisons; *N* = 5-8 per group. **B)** Similar setup as in panel A, but for WT female mice. \**P* < 0.05, \*\**P* < 0.01. Two-way repeated measures ANOVA, F(8, 72) = 4.106; Tukey’s multiple comparisons; *N* = 6-8 per group. **C)** Paw withdrawal threshold in WT male mice before and after SNI surgery, with i.t. AAV injection performed 5 weeks post SNI surgery. \**P* < 0.05, \*\**P* < 0.01, \*\*\**P* < 0.001, \*\*\*\**P* < 0.0001. Two-way repeated measures ANOVA, F(9, 93) = 5.474; Holm-Šidák’s multiple comparisons; *N* = 7-8 per group. **D)** One-year treatment assessment. Paw withdrawal threshold in WT male mice before and after SNI surgery, and with i.t. AAV injection performed 2 days after SNI surgery. \**P* < 0.05, \*\*\**P* < 0.001, \*\*\*\**P* < 0.0001. Two-way repeated measures ANOVA, F(20, 360) = 8.305; Holm-Šidák’s multiple comparisons; *N* = 10 per group. **E)** Paw withdrawal reaction score following brushing of the hind paw before and after SNI or sham surgery with i.t. AAV injection performed 2 days post SNI or sham surgery. Two-way repeated measures ANOVA, F(6, 38) = 9.971, Tukey’s multiple comparisons; *N* = 6 per group. **F)** Latency for first paw withdrawal on a 2°C cold plate before and after SNI or sham surgery with i.t. AAV injection performed 2 days post SNI or sham surgery. Two-way repeated measures ANOVA, F(9, 57) = 13.39, Tukey’s multiple comparisons; *N* = 6 per group. **G)** Total time exhibiting nocifensive behavior during 60 sec following acetone applied to the ipsilateral paw on day 70 with i.t. AAV injection performed 2 days post SNI or sham surgery. One-way repeated measures ANOVA, F(3, 19) = 9.101, Tukey’s multiple comparisons; *N* = 5-6 per group. **H)** Number of spontaneous paw lifts (ipsilateral) recorded over 15 min in SNI or sham surgery mice after i.t. AAV injection. Two-way repeated measures ANOVA, F(3, 19) = 5.013, Tukey’s multiple comparisons; *N* = 5-6 per group. Abbreviations: BL, baseline; i.t., intrathecal. All data in A-H are shown as mean with s.e.m.

### The AAV-encoded PICK1 inhibitor reduces several sensory pain modalities induced by SNI

Neuropathic pain conditions typically manifest within several pain modalities. Therefore, we tested WT male mice receiving AAV8-Di(7P14P)-C5 or AAV8-tdTomato across other pain modalities associated with SNI, now including both SNI and sham groups. Like the 1-year experiment, i.t. injections were performed on day 2 after injury (**Fig. S7A**). In this cohort, we first confirmed by von Frey stimulations that mechanical allodynia was successfully established before the AAV intervention was delivered. In accordance with previous results, injection of AAV8-Di(7P14P)-C5 completely attenuated mechanical allodynia in SNI mice (**Fig. S7B**), while sham mice did not show any change in paw withdrawal threshold (**Fig. S7B**). In the dynamic mechanical allodynia test, using dynamic (brush) instead of static (von Frey hairs) touch, averaged response scores were increased after SNI (compared to sham) and were significantly lower following expression of AAV8-Di(7P14P)-C5 as compared to AAV8-tdTomato at day 40, whereas responses in both sham groups were unchanged (**Fig. 3E**). For cold allodynia tested on a 2°C cold plate, mice subjected to SNI displayed a much lower response latency on day 6 following injury. SNI mice treated with AAV8-Di(7P14P)-C5 showed a significant recovery in their response latency on day 35 and not on day 41 compared to mice injected with AAV8-tdTomato, albeit with considerable variations, which was not observed for the other groups (**Fig. 3F**). Similarly, local paw application of acetone to probe cold allodynia caused aggravated nocifensive behaviors in both SNI groups, but with trending, yet non-significant, treatment differences seen for SNI mice injected with AAV8-Di(7P14P)-C5 and AAV8-tdTomato (**Fig. 3G**). As an index of spontaneous pain, we finally quantified the number of spontaneous paw lifts (SPL) (*24*) for 15 min on day 46 and day 70. While sham mice, regardless of treatments, had almost no SPL, this behavior was prominent in the SNI groups but significantly reduced in mice receiving AAV8-Di(7P14P)-C5 as compared to AAV8-tdTomato (**Fig. 3H**). This suggested that not only provoked but also ongoing spontaneous pain behaviors were attenuated by the treatment. Summarized, this series of behavioral assays support that several sensory pain modalities caused by chronic nerve injury, including dynamic and static tactile allodynia, as well as spontaneous pain behavior, can be attenuated by AAVs encoding recombinant peptides targeting PICK1.

### AAV-encoded PICK1 inhibitors neither improve motor deficits nor alter normal motor functions

Along with the battery of sensory testing, the same SNI and sham groups of mice were temporally exposed to automatic monitoring in single-housed cages to determine potential changes in voluntary behaviors, including rearing, grooming, climbing, immobility, and locomotion. No differences were found between any of the groups (**Fig. 4A, Fig. S7C-I**). Next, we asked whether the therapeutic intervention could reverse any motor function deficits associated with SNI. In CatWalk, SNI mice had significantly lower max contact area and swing speed of the injured paw than sham mice, but no treatment changes were seen across groups (**Fig. S7J-K**). Average run speed and duration in CatWalk were unaffected by SNI and again not affected by treatments across groups (**Fig. S7L-M**). In voluntary wheel running, sham mice ran significantly longer than SNI mice, but no treatment difference was observed within treatment groups (**Fig. S7N)**. This data support that gross motor deficits caused by peripheral nerve transection cannot be attenuated by recombinant PICK1 inhibitors. Apart from this, body weight was monitored periodically over 91 days, with no clear changes seen over time across treatments (**Fig. S7O**). Other phenotypical changes (nest building, social behavior, muscle weakness, etc.) were also not noticed by the experimenters throughout. In summary, the data suggest that general well-being and health status are unaffected by the treatments, but there are noticeable motor deficiencies caused by SNI.

**Fig. 4.**
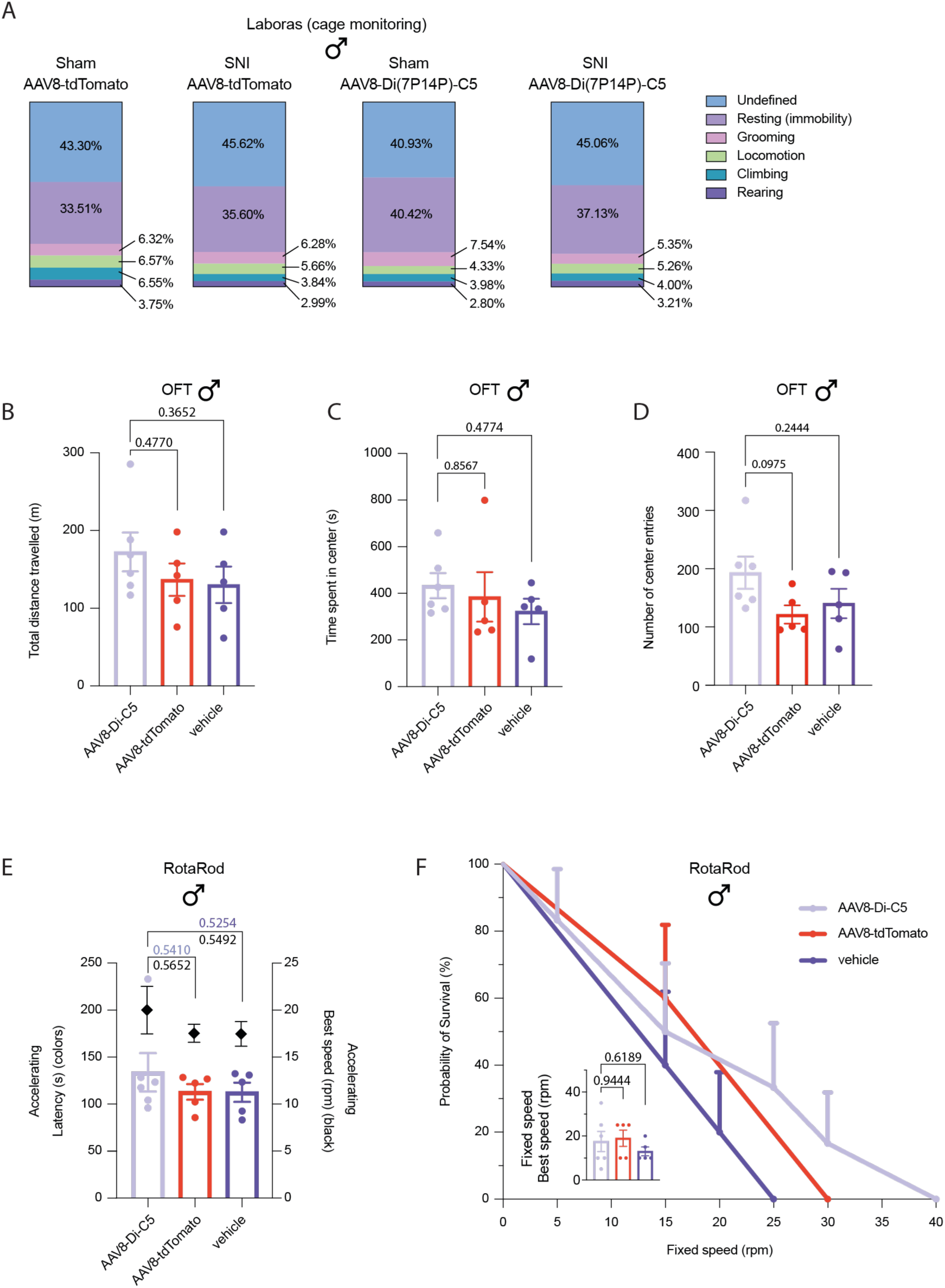
Unperturbed motor function performance following AAV dimeric peptide treatment. **A)** Automatic cage monitoring of general behavior during 22 hours in sham or SNI animals following i.t. injection of AAVs using the Laboras system. Percentages of time spent for each classified behavior are shown. Individual scores and statistics are provided in Suppl. Figure 7C-I. **B)** Total distance traveled in a 2-hour open field test for naïve animals receiving either i.t. injection of AAVs or vehicle (DPBS). One-way ANOVA, F(2, 13) = 0.979; Tukey’s multiple comparisons. *N* = 5-6 per group **C)** Time spent in the center zone in the same test as panel B. One-way ANOVA, F(2, 13) = 0.583; Tukey’s multiple comparisons. **D)** Number of center entries in the same test as panel B. One-way ANOVA, F(2, 13) = 2.459; Tukey’s multiple comparisons. **E)** Same mice as in panel B were examined on the accelerating RotaRod for time to fall (left) and best speed obtained (right). Time to fall: One-way ANOVA, F(2, 13) = 0.671; Tukey’s multiple comparisons. Best speed: One-way ANOVA, F(2, 13) = 0.620; Tukey’s multiple comparisons. Average scores of 3 trials per mouse. **F)** Same mice as in panel B were examined in the fixed-speed RotaRod paradigm. Fall is shown as a survival probability. AAV8-Di-C5 vs. AAV8-tdTomato: Log-rank (Mantel-Cox) test, Chi-square = 0.089, *P* = 0.765; AAV8-Di-C5 vs. vehicle: Log-rank (Mantel-Cox) test, Chi-square = 0.916, *P* = 0.339. Panel insert: Best speed obtained. One-way ANOVA, F(2, 13) = 0.648; Tukey’s multiple comparisons. Abbreviations: i.t., intrathecal; OFT, Open field test. All data in B-F are shown as mean with s.e.m.

We next i.t. injected naïve male mice with AAV8-Di-C5, AAV8-tdTomato, or vehicle (DPBS) to further assess any potential side effects of the AAVs (**Fig. S8A**). In a 2-hour open field test completed 35 days following i.t. injection, total distance traveled, time spent in center, and number of center entries did not differ between treatment groups (**Fig. 4B-D, Fig. S8B**). Mice were also subjected to RotaRod where latency to fall and best speed obtained using accelerating speed remained unaffected across groups (**Fig. 4E**). Likewise, in the RotaRod fixed speed test, no significant difference was found between treatment groups when comparing latency to fall and the best speed obtained (**Fig. 4F**). Together this suggests that the AAV treatments cause no major impact on motor coordination in uninjured mice. We finally verified the efficacy of the AAVs in this setup by subjecting the mice to CFA. Levels of mechanical allodynia indeed confirmed that AAV8-Di-C5, but not by AAV8-tdTomato and vehicle treatment, fully alleviated inflammation-induced mechanical allodynia (**Fig. S8C**), in accordance with previous results (**Fig. 1C**). Together, these results argue that AAV-encoded recombinant peptides targeting PICK1 selectively affect maladaptive pain signaling circuits and processing without alternating gross motor or proprioceptive control in healthy intact circuits.

### Systemic AAV delivery selectively targeting either CNS or PNS completely alleviates neuropathic pain

Novel AAV capsids, like the AAVPHP.S and AAVPHP.eB, can selectively transduce either CNS or PNS, respectively (*25*). Motivated by our previous observations that targeting DRGs seems to play a primary role in suppressing pain by the recombinant peptides delivered by AAV serotype 8, we wanted to test if exclusive peripheral expression would be equally efficacious. AAVs encoding recombinant peptides were packaged into AAVPHP.S or AAVPHP.eB capsids. Mechanical allodynia, repeatedly measured by von Frey stimulation, was confirmed before AAVs were intravenously (i.v.) injected into WT male mice on day 37 after SNI surgery. Control groups receiving either vehicle (DPBS) or AAVPHP.S-iCre:EGFP displayed mechanical allodynia after SNI and did not recover. On the contrary, mice receiving the peptides by PNS (AAVPHP.S-iCre:EGFP-P2A-Di-C5) or CNS (AAVPHP.eB-iCre:EGFP-P2A-Di-C5) targeting capsids recovered completely, as measured up to 125 days after SNI surgery (**Fig. 5A**). Transduction profiles of such AAVs were examined in Ai14 mice receiving identical dosing. The AAVPHP.S capsid gave rise to clear tdTomato labeling of DRG neuronal cell bodies and dorsal column fibers in the spinal cord and brainstem (**Fig. 5B**), almost identical to i.t. injected mice exposed to AAV8 vector (**Fig. 2C-E, Fig. S6C**). A few scattered tdTomato-positive neurons were also observed within CNS without region specificity (**Fig. 5B**). In contrast, the AAVPHP.eB capsid gave rise to strong tdTomato expression in neurons of the spinal cord, brainstem, and brain, while DRG neurons were not labeled (**Fig. 5C**). This pattern was corroborated by the EGFP signal confined towards cell bodies in spinal cord and distributed across the brain (**Fig. 5C**). In summary, the data support that systemic delivery of recombinant peptides targeting either PNS or CNS can fully attenuate chronic neuropathic pain. Still, we cannot rule out whether the scanty CNS neurons transduced by AAVPHP.S could be involved, although this seems unlikely. Interestingly, long-term expression of recombinant peptides did not induce any overt side-effects, as observed by the experimenters, even following extensive CNS exposure provided by AAVPHP.eB. This further supports PICK1 as a compelling pain target and highlights that therapeutic interventions directed towards PICK1 may not only be envisioned as a treatment for chronic neuropathic pain originating from PNS but also CNS. To elaborate this conclusion, we analyzed tissue samples from deceased human donors of both genders. We conducted pull-down experiments, employing the same synthetic biotinylated GCN4-peptides used on mouse tissue. These final experiments confirmed target expression as well as engagement of the targets in human DRG and spinal cord tissue for the PICK1 targeting peptides (**Fig. 5D**). This translational discovery underscores PICK1 as a reachable target for chronic pain treatment with potential clinical significance for AAV therapeutics.

**Fig. 5.**
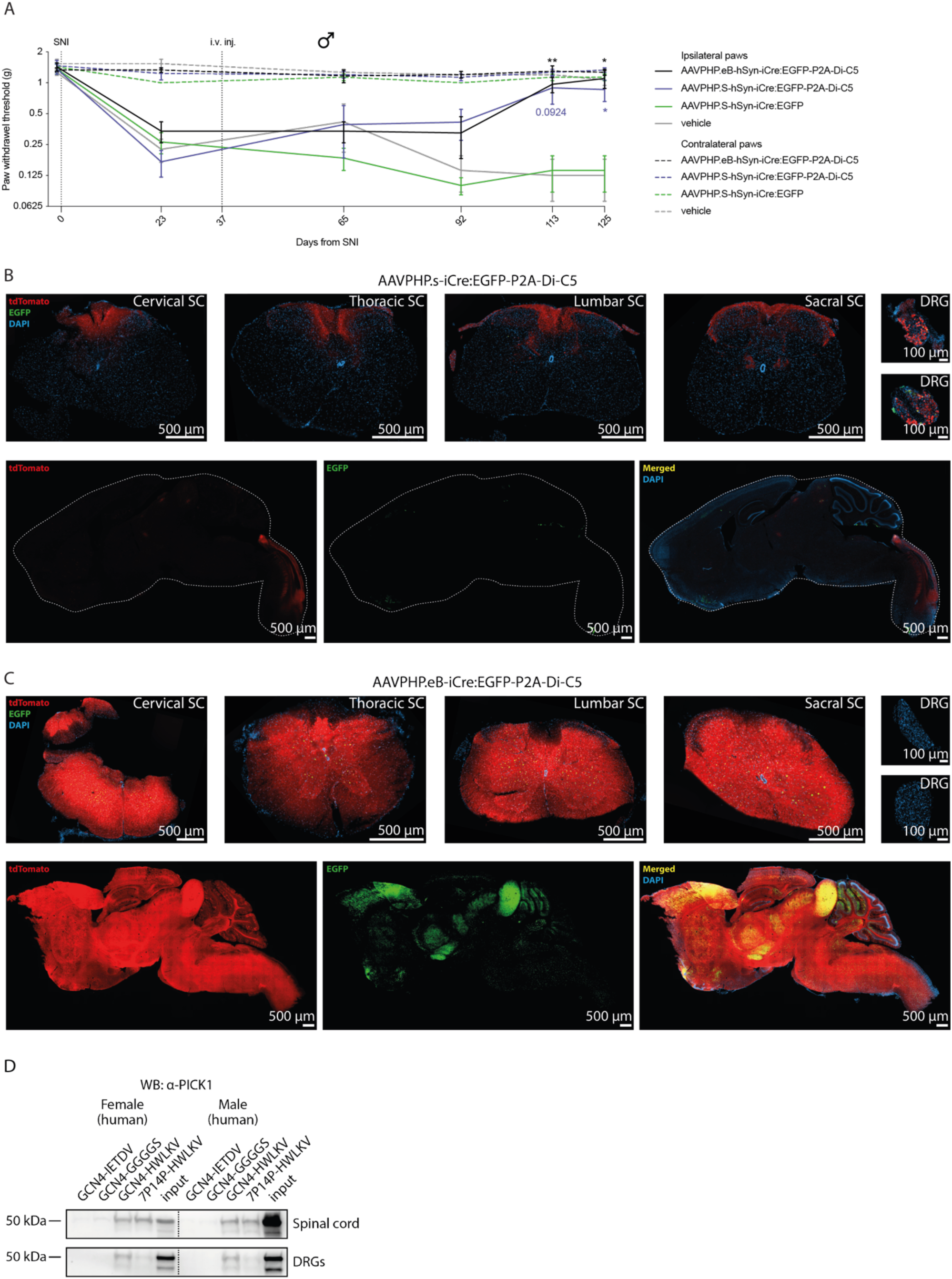
PICK1 inhibitors fully alleviate SNI-induced mechanical allodynia following either CNS or PNS transduction and display target engagement in human tissue. **A)** Paw withdrawal threshold in male WT mice before and after SNI surgery and after systemic AAV administration (i.v. injection) of AAVs with PHP.eB, PHP.S capsids or vehicle (DPBS). \**P* < 0.05, \*\**P* < 0.01. Two-way repeated measures ANOVA, F(5, 100) = 3.727; Dunnett’s multiple comparisons (AAVPHP.S-iCre:EGFP serving as control); *N* = 6 per group. Abbreviations: i.v. = intravenous. Data are shown as mean with s.e.m. **B)** TdTomato signaling from Ai14 mice receiving AAVPHP.S-iCre:EGFP-P2A-Di-C5 vector (*N* = 4) with fluorescent fibers along the spinal cord dorsal column at all levels and with labeled neuronal somas within the DRGs. Ascending fibers terminate at the cuneate and gracile nuclei within the brain stem. Expression of EGFP is visible in DRGs only. **C)** Same as for panel B, but for Ai14 mice receiving AAVPHP.eB-iCre:EGFP-P2A-Di-C5 vector (*N* = 8). TdTomato signaling is not detected in DRGs, but notable throughout the spinal cord and brain. EGFP expression is visible as green dots in the spinal cord and distributed across the brain. **D)** Western blot of PICK1 following pull-down from human spinal cord and DRG tissue homogenate of both genders spiked with biotinylated peptides: GCN4-HWLKV and GCN4(7P14P)-HWLKV peptides target PICK1; GCN4-IETDV targets PSD-95; GCN4-GS is a negative control. *N* = 3 technical replicates for each sample.

## DISCUSSION

Very few treatments for neuropathic pain show long-lasting benefits. Despite the obvious unmet medical need, developing an analgesic with better efficacy and fewer side effects has failed. AAV-based gene therapy is a new therapeutic frontier in medicine and offers the potential to address significant unmet clinical needs (*2, 7*), including the treatment of pain conditions (*26–28*).

In this study, we developed a novel AAV gene therapy, where the pain-relieving effects were confirmed across three different academic institutions. We show that our intervention can reverse neuropathic pain across different sensory modalities by AAV-encoded recombinant dimeric peptides designed to target PICK1 with high affinity. PICK1 is an interesting scaffold protein in the context of pain. It plays a central role in synaptic plasticity and interacts with several proteins of relevance to pain, including several glutamate receptors (GRIK1, GRIK2, GRIK4, GRM7), in addition to the AMPA receptor subunit (GRIA2 and 3), acid-sensing ion channels (ASIC1 and 2), aquaporin (AQP1) and protein kinase C-alpha (PKA-C) (*29*). Furthermore, PICK1 protein is highly conserved and its mRNA is found within neurons of both rodents and humans along the somatosensory nociceptive pathway substantiating that PICK1 is a relevant therapeutic pain target (*29*). While prior research had established the presence of PICK1 protein in mouse DRG neurons (*30*), this study extends this knowledge by demonstrating its expression in human DRG and spinal cord tissue of both genders. Additionally, we provide evidence that the engineered recombinant peptides can selectively bind PICK1 in human tissue.

The therapeutic relevance of PICK1 in the management of persistent pain states is also supported by studies in animal models using inhibitory peptides, siRNA, and knock-out mice, where the lack of PICK1 function was shown to decrease hyperexcitable pain conditions (*30–33*). PICK1 is known as a gatekeeper that controls the balance between calcium-impermeable and CP-AMPARs (*34–36*). Along these lines, a peripherally applied CP-AMPAR antagonist was shown to alleviate chronic pain without affecting mechanosensation or eliciting central side effects (*37*) in concordance with the present data. Nonetheless, due to the very limited insight into PICK1 function in DRGs, we still do not know the significance of other potential PICK1 interaction partners that may contribute to the effect.

The i.t. and i.v. delivery routes in Ai14 mice, utilizing AAV8 and AAVPHP.S, respectively, demonstrated that such methods primarily transduced DRG neurons and their extensions, while neuronal cell bodies of the spinal cord and supraspinal regions remained unaffected. This was further supported by the mRNA peptide profile obtained from WT mice following i.t. injection of AAV8. In accordance, a study with i.t. injection of AAV8-GFP in rats resulted in labeling of the dorsal column fibers and DRG neurons, but with no neuronal cell bodies found outside the DRGs (*38*). This indicates that the therapeutic effect, as seen for AAV8-Di-C5 and AAV8-Di(7P14P)-C5 vectors, may involve ascending dorsal column medial lemniscus (DCML) fibers. This pathway consists of Aβ low-threshold mechanoreceptor (LTMR) fibers that normally transmits non-noxious mechanosensory information. However, nerve injury may cause a phenotypic switch in which Aβ fibers acquire nociceptive capacity, which is usually mediated by small unmyelinated C fibers or thinly myelinated Aδ fibers (*39*). Notably, a study found that tactile allodynia induced by spinal nerve L5/L6 ligation in rats was abolished following complete dorsal column lesion without causing motor deficits (*40*), and another study found that optogenetic activation of Aβ fibers following peripheral nerve injury drove tactile allodynia (*41*). Therefore, pain processing specific to particular modalities by distinct sensory neurons appears to undergo alterations in pathological conditions. So, if the peripheral mechanism of action caused by PICK1 inhibition is equally efficacious in humans, an AAV-mediated gene transfer selectively towards DRGs will be very different from existing therapeutics for the treatment of neuropathic pain and likely also more benevolent.

Before advancing with such AAV treatment for human application, it will be essential to thoroughly address potential translational barriers. While AAV therapeutics are generally well tolerated in humans, the dimeric scaffold, originating from a yeast-derived sequence, may carry a risk of eliciting an immunogenic response. This concern could be mitigated by substituting the dimerization motif with one of human origin. Our findings in the CFA model demonstrate that the dimerization domain can be substituted without compromising efficacy, indicating that alternative, potentially safer motifs can be identified and engineered. Furthermore, while the expression of the dimeric peptide following i.t. delivery of AAV8 predominately targets DRGs, substantial amounts are also detected in the liver, while supraspinal areas are less likely to be transduced. Therefore, to minimize the spread of AAV, a transforaminal epidural injection, used in the clinical practice to deliver steroids for treating lumbar radicular pain, could be advantages for limiting expression towards those DRGs involved in the process and maintenance of peripheral neuropathic pain. Another consideration involves the use of the human synapsin promoter, selected to achieve pan-neuronal expression of the peptides. Surprisingly, whole-mount imaging of Ai14 reporter mice revealed distinct tdTomato signals from what seems to be connective and muscle tissue surrounding the vertebrae. This finding underscores the necessity for thorough evaluation of AAV promoters, even those considered cell-specific, beyond their intended target regions for potential clinical applications.

When packaged with the AAVPHP.eB capsid, transduction and expression of the AAV were restricted exclusively towards neurons in the CNS covering the entire spinal cord and supraspinal areas. Here, equally potent treatment efficacy on mechanical allodynia was seen, as compared to expression in peripheral neurons. This suggests that the recombinant peptides may even be of clinical relevance also for central neuropathic pain conditions. Important limiting factors for clinical AAV gene therapy includes extraordinarily high development and production costs and neutralizing antibodies which limit AAV vectors for repeated administrations. In this study we show long-lasting efficacy for at least 1 year, which of course highlights the long duration of the treatment effect and further argues that the mechanism of action is not prone to desensitization. Taken together, the present work provides evidence that AAV-encoded dimeric peptide inhibitors designed to target PICK1 can effectively provide long-term relief of chronic pain and may represent a promising therapeutic option that can be viewed as a one-shot, life-long treatment.

## MATERIALS AND METHODS

### Experimental Design

Our study aimed to molecularly engineer a class of recombinant peptides expressed by AAVs, designed to form dimers upon translation, that effectively bind PICK1. Furthermore, we sought to assess the efficacy of such dimeric peptides in alleviating sensory pain behavior in mice, while also investigating its expression profile. Additionally, we sought to characterize any potential side effects associated with this treatment and evaluate its ability to target and engage with PICK1 in human-derived tissue samples.

All mice utilized in the study were housed in individually ventilated cages (IVC) and provided *ad libitum* access to water and chow food, under a 12-hour light/dark cycle, and group-housed with 3-8 mice per cage. Prior to the commencement of the experiments, animals were allowed a minimum of 7 days for habituation to the animal facilities. All animal experiments were conducted in AAALAC-accredited facilities and approved by either the Danish Animal Experiments Inspectorate (#2016-15-0201-00976, #2019-15-0201-0160, #2021-15-0201-01036) or by the Regierungspräsidium, Karlsruhe, Germany (#G184/18). Behavioral testing and subsequent data analysis were conducted in a blinded manner with respect to experimental conditions, and each behavioral test was consistently conducted by the same experimenter throughout the study duration. Prior to experiments, animals were randomly assigned to the various experimental conditions, and then tested in the same order on consecutive days. For human tissue sample experimentations, informed consent was obtained from each donor, either via first person consent (a legally binding document) or from the donor’s legal next of kin. This consent meets the standard guidelines concerning informed consent for organ donation for research governed by the United States Uniform Anatomical Gift Act. While no formal power estimation was performed prior to the experiments, the sample sizes were chosen based on previous experience in the field. Further detailed information on Experimental Design, including Materials, is provided as Supplementary Information.

## ACKNOWLEDGMENT

We thank Lone Rosenquist and Nabeela Khadim for excellent technical assistance, and Prof. Ole Kiehn for generous access to transgenic animals. We acknowledge the Core Facility for Integrated Microscopy, Faculty of Health and Medical Sciences, University of Copenhagen.

## Author contribution

GNH cloned all plasmids, produced all AAV vectors, performed most experiments, completed histology, did all data analysis, and made the graphical layouts. KLJ, MR and CBV were involved in SNI experiments. KLJ was involved in SNI and CFA experiments. LJF was involved in CFA experiments and tissue processing. RCB assisted with AAV production, tissue processing, and staining. LS assisted with tissue processing and mouse pull-down experiments. SEJ assisted with tissue processing and performed the immunohistochemistry. RCA and JHL performed qPCR experiments. AHL conducted MD simulations. MBKK performed human pull-down experiments. SPB and GAH did whole-mount tissue clearing procedures and imaging. NRC performed parts of the biochemistry analysis. ATT, CBV, RK, KLM, and ATS supervised the research. GNH, KLM, and ATS conceptualized the study, designed the research, and interpreted data. NRC, KLM, and ATS invented the recombinant dimeric peptides. GNH and ATS wrote the manuscript with contributions from KLM. All authors critically evaluated and approved the final version of the manuscript.

