## Supplementary methods for "Recombinant dimeric PDZ protein inhibitors for long-term relief of chronic pain by AAV therapeutics"

#### SUPPLEMENTARY MATERIALS

##### Material availability

Further information and requests for reagents, plasmids, or AAVs may be directed to and will be fulfilled by the corresponding author.

##### PICK1 and PICK1-A87L protein expression and purification

50 mL LB with kanamycin was inoculated with *Escherichia coli* (BL21-DE3-pLysS) containing a PICK1 or PICK1-A87L encoding plasmid (pET41) and incubated at 37 °C with shaking overnight. The pre-cultures were transferred to 1 L LB media containing kanamycin with continued incubation. When optical density at 600 nm, $OD_{600}$, reached 0.6, protein expression was induced with 1 mM IPTG, and the temperature was turned down to 20 °C with shaking overnight. The cells were harvested through centrifugation (F9 rotor, 7800 x g, 12 min, 4 °C) and suspended in lysis buffer (50 mM Tris, 125 mM NaCl, 2 mM DTT, 1% Triton X-100, 20 µg/mL Dnase 1, 1 tablet of cOmplete™ Protease Inhibitor Cocktail (Roche) pr 200 mL lysis buffer, pH 7.4). After a -80 °C/4 °C freeze-thaw cycle, the proteins were extracted through centrifugation (F20 rotor 36000 x g, 30 min, 4 °C), and the supernatant was incubated with 750 µL PBS-washed Glutathione Sepharose 4B beads (GE Healthcare) for 2 hrs with gentle turning. 3 centrifugation-wash cycles were performed with gentle centrifugation (3000 x g, 5 min, 4 °C) and TBS wash buffer (50 mM Tris, 125 mM NaCl, 2 mM DTT, 0.01% Triton X-100, pH 7.4). The PICK1 or PICK1-A87L beads complex was transferred to a PD 10 gravity column and further washed with 3 column volumes of TBS wash buffer before being collected and incubated with 5 µL thrombin (0.075U/µL stock) overnight at 4 °C with gentle turning. The cleaved samples

were eluted, and absorbance was measured on NanoDrop 2000 at 280 nM. The extinction coefficient of PICK1, $\varepsilon_{A280}$PICK1 = 32320 $M^{-1}*cm^{-1}$. The samples were always kept at 4°C or on ice.

**Synthetic peptides**

The fluorophore-conjugated peptide, 5FAM-di-HWLKV, was conjugated in-house with the fluorophore, 5FAM, through solid-phase peptide synthesis, as described elsewhere (*42*). All other synthetic peptides were purchased from TAG Copenhagen, Denmark, and verified >95% pure through UPLC-MS. These synthetic peptides, if not otherwise stated, were biotin-conjugated through N-terminal Ahx-linkage. Synthesis of biotin-ahx-Atg16-HWLKV and biotin-ahx-MDV1-HWLKV was attempted but unsuccessful due to their complexity. The following peptides (name: primary sequence) were used:

| | |
|---|---|
| HWLKV: | HWLKV |
| 5FAM-di-HWLKV: | 5FAM-$PEG_4$-$(HWLKV)_2$ |
| GCN4-HWLKV: | biotin-ahx-RMKQLEDKVEELLSKNYHLENEVARLKKLVGGGGS-HWLKV |
| GCN4(7P14P)-HWLKV: | biotin-ahx-RMKQLEPKVEELLPKNYHLENEVARLKKLVGGGGS-HWLKV |
| GCN4-GGGGS: | biotin-ahx-RMKQLEDKVEELLSKNYHLENEVARLKKLVGGGGS-GGGGS |
| SSO10a-HWLKV: | biotin-ahx-GEELLEDIRKFNEMRKNMDQLKEKINSVLSIRQGGGGS-HWLKV |
| GCN4-IETDV: | biotin-ahx- RMKQLEDKVEELLSKNYHLENEVARLKKLVGGGGS-IETDV |

**Fluorescence polarization**

Fluorescence polarization (FP) binding saturation assay was carried out with an increasing concentration of recombinant PICK1 or PICK1-A87L and a fixed concentration of synthetic fluorophore-conjugated peptide, 5FAM-di-HWLKV. FP binding competition assay was carried out with a fixed concentration of PICK1 or PICK1-A87L and fluorophore-conjugated peptide, 5FAM-di-HWLKV, and an increasing concentration of an unconjugated test peptide. Both assays were carried out in 96-well plates (Corning, half-area, black, nonbinding) using TBS buffer (1x TBS, 2 mM DTT, 0.01% Triton X-100) and incubated at 4 °C for 20 hrs after mixing to ensure equilibrium was reached. The plates were read on a POLARstar Omega plate reader using an excitation filter at 488 nm and an emission filter at 535 nm. The data were plotted using GraphPad Prism 9.3.1 and fitted to a sigmoidal dose-response curve for the saturation assay or a one-site competition curve for the competition assay. Ki's were automatically calculated by GraphPad Prism using the Cheng–Prusoff equation. The maximum concentration (Cmax) of PICK1 or PICK1-A87L in the saturation assay was 30 µM. The concentration of PICK1 or PICK1-A87L at 75% of maximum binding from saturation assay was used as the fixed PICK1 or PICK1-A87L concentration in the competition assay. The concentration of the fluorophore-conjugated peptide was kept constant at 20 nM for both assays. Cmax of the unconjugated peptide was 100 µM. Three technical replicates were performed simultaneously, and the assay was repeated 2-3 times.

**Fast-protein liquid chromatography**

Fast-protein liquid chromatography (FPLC) was performed with varying ratios of PICK1 to a synthetic peptide in sterile-filtered TBS buffer (1x TBS, 2 mM DTT, 0.01% Triton X-100) on a 24 mL size-exclusion chromatography column (GE Healthcare Superdex™ 200 increase 10/300 GL). Samples were mixed in a total volume of 500 µL in buffer, which was injected into the column, followed by 30 mL buffer. Samples with PICK1:peptide complexes were allowed a minimum of 20 hrs of incubation at 4 °C to reach equilibrium. Samples containing either PICK1 or synthetic peptide were run after a minimum of 20 min incubation on ice. Absorbance was detected at 280 nm.

**Circular dichroism**

For circular dichroism (CD), peptides were diluted to 30 µM in 50 mM NaPi buffer (pH 8.0) and measured on a Jasco J1500 at 25 °C using a quartz cell with a 1 mm pathlength cuvette. Spectra were recorded from 260-190 nm with 0.1 nm step resolution and 50 nm/min scan speed. The mDEG signal was converted to molar ellipticity θ (deg * $cm^2$ * dmol) using the equation θ = (mDEG*106)/(C*N*L), where mDEG is the measured signal, C is protein concentration (µM), N is the peptide residue length, and L is the cuvette pathlength (mm). CD spectra were plotted using GraphPad Prism.

**Molecular dynamics simulation**

Initial structure of GCN4-HWLKV and GCN4(7P14P)-HWLKV were generated in Alphafold (colab v2.1.0) (*43*). Molecular dynamics (MD) simulations were run in GROMACS 2021.4 (*44*) using the Charmm36_ljpme force field(*44*), as this force field best reflected the circular secondary structure of the peptides, as measured with circular dichroism (**Suppl. Figure 1C**). The peptides were aligned with the dimeric structure of the GCN4 basic region leucine zipper (PDB ID: 1YSA) (*45*). The dimers were placed in a cubic box with periodic boundary conditions, with a min 2 nm distance from protein to edge (approx. 11x11x11 nm) and solvated in TIP3P water with 0.1 nM NaCl. The system was energy minimized followed by an equilibration with a constant number of particles, volume, and temperature (NVT) for 100 ps with 2 fs timesteps. The temperature was fixed at 300 K using v-rescale temperature coupling (*46*). This was followed by 100 ps equilibration with a constant number of particles, pressure, and temperature (NPT). The pressure was kept at 1 bar using isotropic Parinello-Rahmen pressure coupling (*47*). One peptide was fixed using a harmonic position restraint with a force constant of 1000 kJ/mol on all atoms. The other peptide was then pushed towards and then pulled away from the first peptide, using GROMACS pull code at a rate of 1 nm per ns and a harmonic potential with a force constant of 500 kJ/mol/$nm^2$. Frames with 0.2 nm distance were extracted from the push and pull trajectories. Each frame was simulated for 10 ns with constant NPT, with the first peptide kept in place by a position restraint and the second peptide fixed using a 500 kJ/mol/$nm^2$ harmonic restraint. The potential of mean force was calculated using the weighted histogram algorithm method (*48*), where the first 100 ps of each window were omitted from the analysis. This process was repeated 5 times for each construct, to obtain mean values and standard errors, which are reported. The free energy of dimerization was estimated as the difference between the minimum free energy value and the bulk value.

**AAV vector design and cloning**

The AAV plasmids (pAAV) were all designed using the same basic template. Transgene expression was driven by the human synapsin 1 (hSyn) promoter for constitutive pan-neuronal expression (*49*). Downstream of the transgene, a woodchuck hepatitis virus posttranscriptional regulatory element (WPRE) and human growth hormone polyadenylation (hGH polyA) was incorporated. The entire gene cassette was flanked by AAV2 inverted terminal repeats (ITR).

*Design of therapeutic transgene and controls*

The therapeutic transgene consisted of the DNA sequence for the basic leucine zipper region of the yeast transcription factor, GCN4-p1 (GCN4) (*50*), a short flexible linker region, and the PICK1-binding motif, HWLKV (*51, 52*). The GCN4(7P14P) variant had two prolines in positions 7 and 14. Alternative zippers were; SSO10a (*53*), Atg16 (*54*) and MDV1 (*55*). The linker region consisted of 4 glycine and 1 serine residues (GGGGS). The extreme C-terminus was composed of HWLKV for achieving PICK1 binding or exchanged with GGGGS for serving as a negative control.

*Tags and accessory elements*

For all pAAVs containing GCN4, GCN4(7P14P), SSO10a, Atg16, or MDV1 sequence, a human influenza hemagglutinin (HA) epitope tag was inserted upstream of the zipper region. This HA-tag was also included in fluorescently tagged transgenes, i.e. EGFP or iCre:EGFP. Posttranslational self-cleavage was obtained by inserting a 2A peptide sequence derived from porcine teschovirus-1 polyprotein (P2A) (*56*) between EGFP/iCre:EGFP and the transgene. To induce expression in Ai14 transgenic mice, codon-optimized Cre (iCre) was fused to EGFP (iCre:EGFP). To achieve Cre-dependent expression with the Cre ON/OFF system, the expression cassette was flanked by two sets of lox sites (loxP and lox2272).

*Cloning of AAV vectors*

DNA sequences of HA-GCN4-GS4-HWLKV, HA-GCN4(7P14P)-GS4-HWLKV, HA-GCN4-GS4-GGGGS, HA-SSO10a-GS4-HWLKV, HA-Atg16-GS4-HWLKV, and HA-MDV1-GS4-HWLKV were ordered in pEX plasmid vectors from Eurofins Genomics, Germany. Restriction sites *Eco53kI* and *EcoRI* were used to excise the transgene of one pEX vector and inserted into pAAV-hSyn-P2A-EYFP-WPREpA excised with *HpaI* and *EcoRI* resulting in pAAV-hSyn-HA-GCN4-GS4-HWLKV. The remaining vectors were made by using the restriction sites *KpnI* and *EcoRI* from pAAV-hSyn-HA-GCN4-GS4-HWLKV and the pEX vectors.

Vectors containing EGFP and/or iCre were cloned from the sequences BsrGI-P2A-HA-GCN4-HWLKV-HindIII and BsrGI-HA-GCN4-GS4-HWLKV-HindIII (ordered from Eurofins Genomics, Germany) cut with *BsrGI* and *HindIII* and inserted into the *BsrGI* and *HindIII*-cut backbones of pAAV-hSyn-EGFP-WPREpA and pAAV-hSyn-iCre:EGFP-WPREpA.

The DIO vectors, pAAV-hSyn-DIO[EGFP-P2A-HA-GCN4-GS4-HWLKV]ON and pAAV-hSyn-DIO[EGFP-P2A-HA-GCN4-GS4-HWLKV]OFF were made using In-Fusion HD Cloning kit (Takara) through PCR amplification of pAAV-hSyn-EGFP-P2A-HA-GCN4-HWLKV with the respective primer sets: 5'-tgctagctcgactagatatccagcacagtggcggcc-3' (Fw) and 5'-ggcgcgcccgccatattgcggccgcttacact-3' (Rv), and 5'-ggcgcgcccgccatagccaccatggtgagcaagggcgaggag-3' (Fw) and 5'-acgaagttatgctagttacactttcagccaatggctgcc-3' (Rv), and inserted into the *NdeI* and *SpeI*-cut or *NdeI* and *NheI*-cut of the pAAV-hSyn-DIO-MSC-WPREpA backbone, respectively.

The vectors pAAV-hSyn-EGFP-WPREpA, pAAV-hSyn-iCre:EGFP-WPREpA, pAAV-hSyn-DIO-MSC-WPREpA , and pAAV-hSyn-TdTomato-WPREpA were already pre-made in the lab. All final AAV vector plasmids were sequenced and validated prior to AAV production.

**AAV production**

Recombinant AAV vectors were produced in-house using a linear polyethylenimine (PEI, MW 25000) triple-transfection protocol and purified with an iodixanol density gradient column. In brief, for each viral construct, 3 x 500 $mm^2$ dishes (Corning) were seeded with 140 million HEK293t cells/dish the day before transfection in DMEM (Glutamax, 10% FBS, 1% P/S, 4.5g/L glucose). The cells were transfected with 400 µg DNA/3 dishes in a 1:4:2 ratio of pAAV : rep/cap : pHelper, and using linear PEI (*57*). The day following transfection, media was exchanged to low serum DMEM (Glutamax, 1% FBS, 1% P/S, 4.5g/L glucose) and incubated for 96 hrs before harvesting of AAV vector in both media and cells using PEG8000 as previously described (*57*). The AAVs were purified from an iodixanol density gradient column following 15 hrs ultracentrifugation (rotor SW28, 28000 rpm, 4 °C). Purity was confirmed with SDS-PAGE, and titer (vg/mL) was determined with Quant-iT PicoGreen dsDNA Assay Kit (Invitrogen). All AAVs are made in serotype AAV2.8 unless otherwise specified. The serotypes AAV2.PHP.S and AAV2.PHP.eB were used for the transduction of PNS or CNS, respectively. All AAV vectors were produced within the lab at University of Copenhagen, and their abbreviated name and full name are as follows:

| | |
|---|---|
| AAV8-Di-C5: | AAV2.8-hSyn-HA-GCN4-GS4-HWLKV |
| AAV8-Di(7P14P)-C5: | AAV2.8-hSyn-HA-GCN4(7P14P)-GS4-HWLKV |
| AAV8-tdTomato: | AAV2.8-hSyn-tdTomato |
| AAV8-Di-SGGGG: | AAV2.8-hSyn-HA-GCN4-GS4-GGGGS |
| AAV8-SSO10a-C5: | AAV2.8-hSyn-HA-SSO10a-GS4-HWLKV |
| AAV8-Atg16-C5: | AAV2.8-hSyn-HA-Atg16-GS4-HWLKV |
| AAV8-MDV1-C5: | AAV2.8-hSyn-HA-MDV1-GS4-HWLKV |
| AAV8-EGFP-P2A-Di-C5: | AAV2.8-hSyn-EGFP-P2A-HA-GCN4-GS4-HWLKV |
| AAV8-EGFP-Di-C5: | AAV2.8-hSyn-EGFP:HA-GCN4-GS4-HWLKV |
| AAV8-iCre:EGFP-P2A-Di-C5: | AAV2.8-hSyn-iCre:EGFP-P2A-HA-GCN4-GS4-HWLKV |

| | |
|---|---|
| AAV8-DIO[EGFP-P2A-Di-C5]ON: | AAV2.8-hSyn-DIO[EGFP-P2A-HA-GCN4-GS4-HWLKV]ON |
| AAV8-DIO[EGFP-P2A-Di-C5]OFF: | AAV2.8-hSyn-DIO[EGFP-P2A-HA-GCN4-GS4-HWLKV]OFF |
| AAVPHP.S-iCre:EGFP-P2A-Di-C5: | AAV2.PHP.S-hSyn-iCre:EGFP-P2A-HA-GCN4-GS4-HWLKV |
| AAVPHP.S-iCre:EGFP: | AAV2.PHP.S-hSyn-iCre:EGFP |
| AAVPHP.eB-iCre:EGFP-P2A-Di-C5: | AAV2.PHP.eB-hSyn-iCre:EGFP-P2A-HA-GCN4-GS4-HWLKV |

**Immunocytochemistry**

Primary cultures of mouse postnatal cortical neurons were transduced with AAVs and incubated for a minimum of 6 days. The cells were fixed, permeabilized, and blocked (5% serum, 0.25% triton-X 100), and incubated with rabbit α-GCN4 (1:275, C11L34, Absolute Antibodies) and A488 goat α-rabbit (1:5000, A11034, Invitrogen) before stained with mounting media containing DAPI. Cells were imaged with a Zeiss LSM 780 laser scanning confocal microscope with a 63x objective. Images were processed using ImageJ or Zen (Zeiss software).

**Animals**

Unless otherwise specified, male or female C57BL/6NRj, male C57BL/6JRj (Janvier, France), or male C57BL/6JBomTac (Taconic, Denmark) wild-type (WT) mice between 7-12 weeks of age were ordered externally. The Ai14 (JAX #:007908) and Hoxb8-Cre$^{+/-}$ (JAX #:035978) mice strains were generously provided by Prof. Ole Kiehn and propagated within the animal facilities. Homozygotes Ai14$^{+/+}$ and heterozygous Hoxb8-Cre$^{+/-}$ mice of both sexes were used for experiments. Mice were housed in individually ventilated cages (IVC) with access to water and chow food *ad libitum*, maintained on a 12-hour light/dark cycle, and group-housed (3-8 mice per cage). Animals were allowed at least 7 days of habituation to the animal facility before the initiation of the experiment. Within cages, all mice were exposed to the same AAV and/or surgery, except for CFA experiments. All experiments involving animals were performed in AAALAC-accredited animal facilities and approved by the Danish Animal Experiments Inspectorate (#2016-15-0201-

00976, #2019-15-0201-0160, #2021-15-0201-01036) or by the Regierungspräsidium, Karlsruhe, Germany (#G184/18).

**Pull-down and western blot (mouse)**

*Lysate preparation.* DRGs L1-L5, lumbar spinal cord, and brainstem located beneath cerebellum were dissected and pooled from 2 naïve male WT mice and immediately homogenized in 400 µL ice cold lysis buffer (100 mM NaCl, 50 mM Tris-Cl, 1% NaDeoxycholate, protease inhibitor cocktail (10 µL in 1 mL lysis buffer, P8340, Sigma) followed by agitation on spinning wheel (4 °C, 30 min). The lysates were centrifugated (4 °C, 13.000 x g, 15 min) and the supernatant collected.

*Pull-down.* The mouse lysates were pre-cleared with 3x lysis buffer-washed streptavidin beads (15µL, Dynabeads™ MyOne™ Streptavidin T1, 65601, Invitrogen) on a spinning wheel (4 °C, 1 hr). Total protein in the lysates was determined by a BCA assay (Thermo Scientific) and read on Omega POLARstar plate reader (BMG Labtech). Simultaneously, streptavidin beads (30 µL) were pre-incubated with 10 µM biotinylated peptide (GCN4-GGGGS, GCN4-HWLKV, GCN4(7P14P)-HWLKV, or GCN4-IETDV) in 500 µL TBS buffer (1x TBS, 2 mM DTT, 0.01% Triton X-100) on a spinning wheel (4 °C, 3 hrs), followed by removal of unbound peptide through 3x washes in TBS buffer. Pre-cleared lysate (500 µg protein) was added to pre-incubated bead-peptide mix and incubated overnight on spinning wheel (4 °C). Unbound material was gently removed through 3x washes in TBS buffer.

*SDS-PAGE and western blotting*. The pulled peptides/proteins were eluted from the beads with 30 µL 2x Laemmli buffer and boiling (100 °C, 6 min). Input (30 µg protein) was likewise boiled in 30 µL 2x Laemmli buffer. The resulting pulled peptides/proteins and input were run on a 4-15% gradient SDS-PAGE gel (#4561083, Bio-Rad) and transferred to a membrane. The blot was blocked in 5% (w/v) non-fat dry milk (9999, Cell Signaling Technology) in PBS-T buffer (PBS with 0.01% Tween 20, P9416, Sigma-Aldrich) and incubated in primary antibody against PICK1 (1:1000, mouse monoclonal #75-040, NeuroMab) or PSD-95 (1:500, mouse monoclonal, ab192757, Abcam) (4 °C, overnight, gentle shaking). The blot was washed 5 x 5 min in PBS-T buffer and incubated in secondary antibody (1:2000, Goat anti mouse IgG HRP, #31430, Pierce) in PBS-T buffer (1.5 hrs, gentle shaking) followed by 5 x 5 min washes in PBS-T buffer. The protein complexes were analyzed on Alpha Innotech (Flour H2D software) with SuperSignal ELISA Femto Substrate (#37075, Thermo Scientific) for PICK1 and ECL Prime Western Blotting Detection (#89168-782, GE Healthcare) for PSD-95.

**Human donor tissue**

Two DRG pairs from the lumbar region (L3-L5) from one female (46 years, Caucasian) and one male (52 years, Caucasian) and two spinal cord sections from the lumbar region (L3-L5) from one female (52 years, Caucasian) and one male (55 years, Caucasian) human donors were acquired from AnaBios, San Diego, US. None of the donors had any known neuropathic pain history. Tissues were flash-frozen in liquid nitrogen upon dissection and shipped in temperature-controlled conditions. Informed consent was obtained from each donor by AnaBios, specifying that the donor tissue can be utilized strictly for laboratory purposes only.

**Pulldown and western blot (human)**

Lysate preparation, pulldown, SDS-PAGE, and western blotting were performed as done for mouse with the following exceptions.

*Lysate preparation:* Lysates was made from DRGs and spinal cord sections from 2 female and 2 male human donors.

*Pulldown:* 50 µg protein was used for pulldown conditions and 20 µg protein for input.

*SDS-PAGE and western blotting*: Proteins were eluted with 20 µL Laemmli buffer, and an Any kD SDS-PAGE gel (#4569035, Bio-Rad) was used. The protein complexes were analyzed on Amersham ImageQuant 800 (Cytiva) with SuperSignal ELISA Femto Substrate (#37075, Thermo Scientific).

**Immunohistochemistry and imaging of slice tissue**

*Cryosection samples*

Ai14 reporter mice receiving 7 µL i.t injection of AAV2.8-iCre:EGFP-P2A-Di-C5 mice were perfused with PBS followed by 4% paraformaldehyde (PFA) 40-44 days post injection. Brain, spinal cord, and DRGs were dissected, and tissue was kept in 4% PFA overnight, stored in 30% sucrose for 2 days, and finally embedded in OCT (TissueTek). DRGs (10 µm), spinal cords (20 µm), and brains (40 µm) were cut a cryostat (Leica CM3050 S). Tissue sections were mounted with media containing DAPI and imaged using AxioScan Z1 with 20x objective. Image visualization was performed using Zen (Zeiss) and Fiji software.

For DRG immunohistochemistry, 10 µm DRG sections were blocked in blocking buffer (PBS with 0.3% Triton X-100 and 10% donkey serum) for at least 1 hr before they were incubated in primary antibodies diluted in blocking buffer overnight at room temperature. Next, they were washed in PBS with 0.3% Triton x-100 3x5 min, incubated in secondary antibodies diluted in blocking buffer for at least 4 hrs, washed 3x5 min and mounted with mounting medium. The stained DRG sections were imaged on a Zeiss Axio Scan.Z1 slide scanner and analyzed using Zeiss Zen software. Primary and secondary antibodies, including their dilution, used:

tdTomato: Rabbit anti-mCherry (1:200, PA5-34974, Invitrogen); Goat anti-rabbit 568 (1:500, A11036, Invitrogen) or Donkey anti-rabbit 568 (1:500, A10042, Life technologies).

IB4: IB4-647 (1:500, I32450, ThermoFisher).

CGRP: Goat anti-CGRP (1:400, ab36001, Abcam); Donkey anti-goat 647 (1:500, A21447, Invitrogen).

NF200: Chicken anti-NF200 (1:500, Ab5539, Millipore); Goat anti-chicken 647 (1:500, A21449 Life technologies).

**Whole-mount tissue clearing**

*Fixation protocol*

Ai14 mice received a 7 µL intrathecal injection of AAV2.8-iCre:EGFP-P2A-Di-C5. Forty days post-injection, mice were euthanized and perfused with 20 mL of PBS followed by 20 mL of 4% PFA to ensure fixation. A tissue sample was dissected with the entire vertebrae and skull attached. Samples were post-fixed in 4% PFA for 24 hours at 4°C, then washed with PBS containing sodium azide (NaN3) 0.02%.

*Tissue-Clearing and immunolabeling*

The tissue-clearing and immunolabeling processes were adapted from the iDISCO protocol (*58*) with several modifications to enhance tissue preparation and labeling efficiency. Initially, decalcification was conducted by immersing the samples in 20% EDTA (ED4SS, Sigma-Aldrich) at a pH of approximately 7.4, maintained at 37°C for four days. Following decalcification, the samples were thoroughly washed with water and dissected to remove the skull, preparing them for subsequent processing.

Before initiating the standard iDISCO protocol, we incorporated an additional delipidation and bleaching phase. This involved treating the samples with 25% Quadrol (#122262, Sigma-Aldrich) for 48 hours at 37°C, followed by 5% ammonium hydroxide (#35574, ThermoFischer) for 24 hours at the same temperature. After these preparatory steps, the samples were processed according to the iDISCO method, including the standard immunolabeling procedures. The primary RFP antibody was incubated for 10 days (1:1000, #600-401-379, Rockland) and the 647 donkey anti-rabbit (1:1000, #711-605-152, Jackson ImmunoResearch) secondary antibody was incubated for 7 days at 37°C. Following the application of the secondary antibody, an extra post-fixation step was introduced to stabilize the immunocomplexes, involving overnight incubation in 2% PFA at 4°C. This additional step ensured enhanced preservation of tissue morphology and labeling fidelity prior to the final dehydration process. Ethyl cinnamate (W243000, Sigma-Aldrich) was used as the final clearing solvent following the two washes of dichloromethane.

**Light Sheet Microscopy**

The acquisitions were done on a Zeiss LS7 scanning Gaussian-beam light sheet microscope with orthogonal light paths for illumination and detection. Two 5x illumination objectives (Carl Zeiss, NA 0.1) were used to generate dual side illumination of the sample, together with an EC Plan-NEO 5 x detection objective (Carl Ziess, NA 0.16, 10.5 mm working distance). Images were acquired with two PCO Edge 4.2 M sCMOS cameras using Zeiss Zen Black software. The laser line used to capture the autofluorescence signal in the sample was 488 nm (Diode laser, 30 mW). The laser line used to capture the RFP signal was 638 nm (Diode laser, 75 mW). To refine the emission signal additional filters were used, i.e. BP (band pass) 505-545 and LP (long pass) 660 for the corresponding excitation.

**qPCR**

Relative gene expression was examined through quantitative Real-Time PCR (qRT-PCR). Seven weeks old WT animals (n=4) were i.t. injected with AAV-Di-C5 (7 µL of 2.2E12 vg/mL; total viral load 1.5E10 vg/mouse), or 7 µL of DPBS. Four weeks later the mice were sacrificed and sciatic nerve, DRGs (L3-L5 from both sides), lumbar spinal cord, dorsal column, cervical spinal cord, pons, cerebellum, and liver were dissected out. Freshly dissected tissue was immediately flash-frozen on dry ice. Hereafter, tissue samples were stored at -80 degrees until RNA extraction. All surfaces and tools were pre-treated with 70% ethanol and/or RNase*Zap*® Solution (ThermoFisher) to prevent RNAse activity. Each tissue sample was directly placed into 1ml of QIAzol Lysis Reagent (Qiagen), followed by homogenization with a handheld homogenizing pestle. Complete lysis and RNA extraction were performed as previously described (*59*). Reverse transcription was performed with Maxima First Strand cDNA Synthesis Kit (ThermoFisher) using a total of 500 ng RNA from each sample or 200 ng RNA from the sciatic nerve. The resulting complementary DNA (20ml reaction) was further diluted with 125 ml EB buffer (Qiagen). Real-time quantitative PCR was performed on a LightCycler 480 II instrument (Roche Life Science), as previously described (*59*). A specific primer for AAV-Di-C5 mRNA was designed and validated prior to the experiment, using both transduced cell culture samples and non-transduced tissue samples. Housekeeping genes *Eef1a1* and *Yhwaz* are described elsewhere (*59, 60*). The primer efficiencies were ascertained through a seven-step dilution curve. The corresponding Ct value for each well was calculated using the on-board software (Roche) with a maximum cut-off of 35 cycles. For each independent experiment, samples were run in technical triplicates and averaged Ct values were used for calculations (*61*).

Equation 1: $\Delta Ct = mean\ Ct_{gene\ of\ interest} - mean\ Ct_{housekeeping\ genes}$

Equation 2: $\Delta\Delta Ct = mean\ \Delta Ct_{DPBS\ group} - mean\ \Delta Ct_{gene\ of\ interest}$

The following primer pairs were designed:

Eef1a1 (forward): TGCTGGAGCCAAGTGCTA AT; Eef1a1 (reverse): GTGCCAATGCCGCCAATTTT

Yhwaz (forward): GAAGCATTGGGGATCAAGAA; Yhwaz (reverse): AGACGGAAGGTGCTGAGAAA

AAV-Di-C5 (forward): ATCCGTATGATGTGCCGGATT; AAV-Di-C5 (reverse): CGCCACTTCGTTTTCCAGATG

**Pain models**

*Complete Freund's Adjuvant (CFA) model of inflammatory pain.*

Mice were briefly anesthetized with 2% isoflurane (max. 60 seconds) and unilaterally injected in the plantar surface of the right hind paw with 50 µL undiluted Complete Freund's Adjuvant (CFA) (F5881, Sigma) with a single intraplantar (i.pl.) injection using an insulin needle (0.3 mL BD Micro-Fine). Analgesic treatment was not applied to CFA-injected animals.

*Spared nerve injury (SNI) model of neuropathic pain and sham surgery*.

Briefly, mice were deeply anesthetized with 2% isoflurane, and unilaterally a small incision was made in the skin of the left hind leg between the thigh bone and knee joint. Using blunt dissection, the muscles were separated to expose the sciatic trifurcation point. The tibial and common peroneal nerve branches were ligated with a surgical knot and a small piece of the nerve was cut out distally to the ligation which was left untouched. The muscles were tucked into place, the skin was closed with glue and/or sutures, and the health status of the mice was carefully monitored, especially in the immediate days following the surgery. Sham surgeries were performed similarly, exposing the sciatic trifurcation point, but without ligation of the nerve branches. Differences in surgical and analgesic procedures between study sites are highlighted below:

*University of Copenhagen*: The tibial and peroneal nerve branches were ligated with a single surgical knot (nonabsorbable polypropylene 6-0 sutures) (8660H, Ethicon). The skin was closed with glue (Tissue adhesive, Klinibond) (115 500, Klinion). Buprenorphine (Temgesic; 0.3 mg/ml, RB Pharmaceuticals, diluted in isotonic saline from Fresenius Kabi) was administered s.c. 5 min before surgery. A lidocaine/bupivacaine mixture (2.5 mg/mL / 1.25 mg/mL) was topically applied to muscle layer during surgery. Rimadyl (Carprofen, 1% from 0.5 mg/mL stock) was injected s.c. after surgery. Buprenorhpine and Rimadyl treatment was repeated once daily for another 2 days.

*Aarhus University*: The tibial and peroneal nerve branches were ligated with a single surgical knot (nonabsorbable polypropylene 6-0 sutures) (8660H, Ethicon). The skin was closed with glue (Tissue adhesive, Klinibond) (115 500, Klinion). Lidocaine SAD (10 mg/ml; Amgros I/S) was applied once to the skin. Buprenorphine (Temgesic, 0.3 mg/ml; RB Pharmaceuticals) and ampicillin (Pentrexyl, 250 mg/ml; Bristol-Myers Squibb) were mixed and diluted 1:10 in isotonic saline (9 mg/ml; Fresenius Kabi), and 0.1 ml was injected s.c. following the surgery. Buprenorphine and Pentrexyl treatment were repeated once daily for another 2 days, and always given subsequent to behavioral assessments, if performed on Day 2.

*Heidelberg University*: The tibial and peroneal nerve branches were ligated with a single surgical knot (absorbable 5-0 sutures) (17241041, Catgut GmbH). The skin was closed with several surgical knots (absorbable 5-0 sutures) (17241041, Catgut GmbH) followed by glue (1050052, B. Braun Surgical). Only isoflurane anesthesia was used.

**AAV administration**

AAVs were administered either by intrathecal (i.t.) or intravenous (i.v.) injection. For i.t. injections, mice were anesthetized with 2% isoflurane (max. 2 min). Using a hand-held Hamilton syringe (30G, 22mm needle) inserted at the L5/L6 intervertebral space, 7 µL AAV (2.2-3.3 x $10^{12}$ vg/mL; total viral load 1.5-2.3 x $10^{10}$ vg/mouse) was injected once a Straub tail response was elicited. For i.v. injections, mice were placed in a warming chamber to allow the tail vein to swell, and 50 µL (PHP.S, 2.0 x $10^{13}$ vg/mL, total viral load 1.0 x $10^{12}$ vg/mouse; PHP.eB, 2.0 x $10^{12}$ vg/mL, total viral load 1.0 x $10^{11}$ vg/mouse) was injected using an insulin syringe.

**Behavioral tests**

Behavioral testing and following data analysis were performed blinded to experimental conditions, e.g., without knowing which AAVs were used. In general, prior to experimental onset, mice were allowed to habituate to the room for a minimum of 30 min or until mice were calm. Individual experiments were performed by the same female experimenter throughout the duration of the experiment.

*Von Frey*

Von Frey was performed at three different study sites. In common, all sites used the ascending von Frey method with manual von Frey filaments (Bioseb) ranging from 0.04-2.0g (0.04, 0.07, 0.16, 0.4, 0.6, 1.0, 1.4, 2.0g). Each filament was applied to the lateral plantar surface of each hind paw 5 times. Positive responses were used to determine the paw withdrawal threshold (PWT).

*University of Copenhagen*: Mice were placed on an elevated wire mesh in 8 (Ø) x 7.5 (h) cm red, transparent, plastic cylinders or in 11.5 (w) x 14 (d) cm PVC plastic boxes and allowed to habituate for a minimum of 20 min or until calm before the initiation of the experiment. Each filament was applied 5 times to the hind paw of the same mouse over approx. 30-60 seconds or longer resting periods if required. The paw withdrawal threshold was determined as 3 positive responses out of 5 for two consecutive filaments. A positive response was defined as a sudden paw withdrawal, flinching, and/or paw licking induced by the filament.

*Aarhus University*: Mice were placed on an elevated wire mesh in 8 (Ø) x 7.5 (h) cm red, transparent, plastic cylinders and allowed to habituate for a minimum of 15 min before the initiation of the experiment. Each filament was applied 5 times to the hind paw of the same mouse over 30 seconds (each stimulus lasting approx. 2 seconds), followed by application of the filament to the contralateral hind paw. The paw withdrawal threshold was determined as 3 positive responses out of 5 for the same filament. A positive response was defined as a sudden paw withdrawal, sudden flinching, or sudden paw licking.

*University of Heidelberg*: Mice were placed on an elevated wire mesh in 9.5 (w) x 9.5 (d) + 13.5 (h) cm plastic boxes and allowed to habituate for a minimum of 30 min or until calm before the initiation of the experiment. Each filament was applied to each hind paw of each mouse over 5 rounds, giving a total of 5 applications to each hind paw with a minimum of 5 min resting period between each application of the same filament. Between different filaments a resting period of minimum 5 min resting period was used between each filament size. An additional resting period of 5 min was used between each filament size. The paw withdrawal threshold was determined as 3 positive responses out of 5 for two consecutive filaments. A positive response was defined as a sudden paw withdrawal, flinching, and/or paw licking induced by the filament.

*Brush*

Mice were placed on an elevated metal grid in 9.5 (w) x 9.5 (d) + 13.5 (h) cm chambers with lids, and left to habituate for 30 min. A soft, round, size 3 brush was applied in one dynamic motion in a heel-to-toe direction and the paw withdrawal reaction was scored as follows: 0 = no response, 1 = brief lifting, 2 = paw lift and guarding, 3 = paw flinch, 4 = paw flick or multiple paw flinches, 5 = paw licking, or flick and guarding. The brush was applied a total of 3 times for each paw. Data are shown as the average score of 3 applications.

*Acetone test*

Mice were placed on an elevated metal grid in 9.5 (w) x 9.5 (d) + 13.5 (h) cm chambers with lids, and left to habituate for 30 min. A single drop of acetone was applied to the ipsilateral paw with a syringe with a pipette tip. A camera recorded the behavior for 1 min, and the videos were manually assessed for the duration of the nocifensive behavior of the ipsilateral paw, including paw flick, licking, and guarding.

*Coldplate*

The mice were placed on a 2 °C cold plate (Bioseb) and the latency for the first paw withdrawal was recorded. A cutoff of 30 sec was used.

*Dynamic hotplate*

For the dynamic hotplate (Bioseb), the mice were placed on a hotplate with a temperature ramp starting at 30 °C (± 0.1) and rising at 5 °C/min increments until the first nocifensive response (paw withdrawal, flick, or lick). A cutoff temperature was set to 50°C. The data show the temperature at which the nocifensive response occurred.

*Spontaneous paw lifting (SPL)*

Mice were placed in a 19 (w) x 19 (d) x 13.5 (h) cm box with 9.5 (w) x 9.5 (d) cm chambers. The box was placed on a transparent surface allowing a video camera to record the mice from below. Mice were recorded for 15 min. The videos were manually assessed for ipsilateral paw lifting in absence of exploratory behavior including rearing, walking, grooming, and coordinated movements of other paws. Only paw lifts, when the mouse was standing still and no other paws were coordinately lifted, were counted.

*Voluntary wheel running*

Mice were placed in individual cages containing a wheel with free access to food and water. The number of wheel rotations was collected with AWM counter and AWM software (Lafayette Instruments, Louisiana, USA), using optical sensors to detect revolutions. 1 rotation equals 0.4 meters. The mice entered the cages during the light phase for habituation, and data were collected in 1-hour intervals during the dark phase overnight (18:00-06:00). The data represent 3 technical replicates.

*Cage monitoring (Laboras)*

Mice were placed in individual Laboras-specified cages (Metris B.V., Netherlands) placed on a carbon fiber platform that detects a range of behaviors based on the vibration pattern, which was collected with the Laboras 2.6 software. Behaviors assessed consist of frequency and time courses of grooming, rearing, climbing, locomotion, and immobility. Mice were assessed over a 22-hour period and had free access to food and water.

*Open field test*

Mice were placed in the center of square white open arenas (40 x 40 x 40 cm) and monitored for 120 min. Open-field locomotion was recorded and analyzed (distance and position in the arena) using Ethovision video-tracking software (Noldus) with distance traveled further divided into 5 min bins.

*CatWalk*

Mice were placed on the CatWalk XT (Noldus) and allowed 3 runs with the following criteria: average run speed < 30 cm/s and run variation < 100. The data shown represent an average of 3 runs. The post-run analysis included the following parameters: run duration (s), average speed (cm/s), the maximum contact area of the ipsilateral hind paw (LH_MaxContactArea) (cm2), swing speed of the ipsilateral hind paw (LH_SwingSpeed) (cm/s). CatWalk settings during run: Camera Gain = 20.7 dB, Green Intensity Threshold = 0.12, Red Ceiling Light = 17.7 v, Green Walkway Light = 16.5 v.

*RotaRod*

RotaRod was performed over 3 days. On day 1, the mice were trained on the RotaRod. Once 3 min at 5 rpm was completed for each mouse, a 3-days accelerating paradigm was started. Over 5 min, the speed was increased from 4-40 rpm (1 rpm/8 sec). 3 trials per mouse were performed on day 1-3. On day 3 following the accelerating paradigm, the mice were tested with a fixed speed paradigm. The mice were placed on the rod at 5, 10, 15, 20, 25, 30, 35, and 40 rpm for 5 min each time, or until the mouse fell off 3 times at the same speed. An inter-trial break of at least 10 min was used between each speed.

*Study sites for behavioral assessments*

The behavioral experiments were performed by female experimenters at three different study sites: University of Copenhagen (Denmark), Aarhus University (Denmark), and Heidelberg University (Germany). The below list displays the study site where the behavioral experiments were performed.

| | |
|---|---|
| Figure 1C + 1F: | University of Copenhagen |
| Figure 3A, C-D: | Aarhus University |
| Figure 3B: | University of Copenhagen |
| Figure 3E-H: | Heidelberg University |
| Figure 4A: | Heidelberg University |

| | |
|---|---|
| Figure 4B-F: | University of Copenhagen |
| Figure 5A: | University of Copenhagen |
| Suppl. Figure 3A-B: | University of Copenhagen |
| Suppl. Figure 5C-D: | University of Copenhagen |
| Suppl. Figure 6D: | University of Copenhagen |
| Suppl. Figure 7B-O: | Heidelberg University |
| Suppl. Figure 8B-C: | Heidelberg University |

**Statistical analysis**

Statistical details are reported in the figure legends and were computed with GraphPad Prism 9. One-way ANOVA was used to compare group means between more than two groups, followed by either Dunnett's multiple comparisons where groups were compared with controls or Tukey's multiple comparisons where means of every two groups were compared. Two-way ANOVA was used to compare group means when there were two or more variables, followed by either Tukey's, Holm-Šidák's, or Dunnett's multiple comparisons (as noted in the figure legends). Depending on the experimental design, either two-way ANOVA or two-way repeated measures ANOVA was used. One sample t-test was used to compare group means. Survival probability was determined by Log-rank (Mantel-Cox) test, whereas cumulative distribution was determined by Kolmogorov-Smirnov (KS) test. All comparisons are two-sided. Statistical significance was determined as follows: *$P < 0.05$, **$P < 0.01$, ***$P < 0.001$, ****$P < 0.0001$, as indicated in figures where von Frey filaments were applied. Results show mean with s.e.m, if not otherwise stated. No estimate of power was performed before experiments, but sample size numbers were similar to those generally employed in the field.

#### SUPPLEMENTARY FIGURES

**Fig. S1. Self-assembly and helicity are more pronounced for GCN4-HWLKV than for GCN4(7P14P)-HWLKV peptides.**

**A)** Molecular dynamics (MD) simulation for estimation of the free energy of dimerization, which were 84.4 ± 3.7 kJ/mol for GCN4-HWLKV (purple) and 60.9 ± 3.4 kJ/mol for GCN4(7P14P)-HWLKV (lavender). The free energy was shown as mean with s.e.m. from $N$ = 5 simulations per peptide.

**B)** Fast-protein liquid chromatography (FPLC) trace of GCN4-HWLKV (purple) and GCN4(7P14P)-HWLKV (lavender) peptides (10 µM) for assessment of their oligomeric state.

**C)** Circular dichroism (CD) spectra of GCN4-HWLKV (purple) and GCN4(7P14P)-HWLKV (lavender) peptides (30 µM) for assessment of their secondary structure.

**Fig. S2. PICK1 binding and oligomerization are more pronounced for GCN4-HWLKV than for GCN4(7P14P)-HWLKV peptides.**

**A)** Fluorescence polarization (FP) competition binding curves for GCN4-HWLKV (purple), GCN4(7P14P)-HWLKV (lavender), and HWLKV (gray) peptides using a constant concentration of PICK1 and 5FAM-di-HWLKV (20 nM) as a tracer. Ki values from the binding curves are shown to the right, $N$ = 9 (3 separate experiments, each with 3 technical replicates).

**B-E)** Fast-protein liquid chromatography (FPLC) traces of PICK1 **B)** alone (cyan) (40 µM), **C)** with GCN4-HWLKV (lavender) (molar ratio 1:4), **D)** GCN4(7P14P)-HWLKV (purple) (molar ratio 1:4) and **E)** with GCN4-HWLKV (lavender) (molar ratio 1:2). Gray columns indicate elution volume of the peptides alone from Fig. S1B.

**Fig S3. AAVs with EGFP fusion cassettes provide pain relief in the CFA model.**

**A)** Paw withdrawal threshold measured by von Frey stimulation before and after i.t. injection of AAVs, and again after CFA and NTX administration as indicated. Paw withdrawal threshold (ipsilateral) comparison was made between AAV8-tdTomato (control) and the two EGFP-containing vectors, including AAV8-Di-C5 and AAV8-Di(7P14P)-C5 from Figure 1C, as testing of these vectors was completed within the same experiment. *$P < 0.05$, **$P < 0.01$, ***$P < 0.001$, ****$P < 0.0001$. Two-way repeated measures ANOVA, $F(32, 208) = 2.198$; Dunnett's multiple comparisons; $N$ = 5-8 per group.

**B)** Temperature reached for first nocifensive reaction on the dynamic hot plate in CFA and naïve mice, i.t. injected with AAV8-tdTomato or AAV8-Di-C5. One-way ANOVA, $F(3, 36) = 4.971$; Tukey's multiple comparisons; $N$ = 8-10 per group.

Abbreviations: CFA, Complete Freund's Adjuvant; BL, baseline; i.t., intrathecal; NTX, naltrexone. All data in A-B are shown as mean with s.e.m.

**Fig. S4. Substitution of the PDZ binding sequence abolishes PICK1 binding and oligomerization, whereas substitution of the dimerization motifs is tolerated.**

**A)** Fluorescence polarization (FP) competition binding curves for GCN4-HWLKV (lavender), SSO10a-HWLKV (orange), GCN4-GGGGS (gray), and HWLKV (black) peptides using a constant concentration of PICK1 and 5FAM-di-HWLKV (20 nM) as tracer. Relative affinities are shown to the right, $N$ = 2.

**B)** Fast-protein liquid chromatography (FPLC) traces of GCN4-HWLKV (lavender), SSO10a-HWLKV (orange), and GCN4-GGGGS (gray) peptides (20 µM) for assessment of their oligomeric state.

**C)** Circular dichroism (CD) spectra of SSO10a-HWLKV (orange) and GCN4-GGGGS (gray) peptides (30 µM) for assessment of their secondary structure.

**D-F)** Fast-protein liquid chromatography (FPLC) traces of **D)** PICK1 (cyan) (40 µM) with GCN4-HWLKV (lavender) (molar ratio 1:2), **E)** PICK1 (cyan) (40 µM) with SSO10a-HWLKV (orange) (molar ratio 1:2) and **F)**

PICK1 (cyan) (40 µM) with GCN4-GGGGS (gray) (molar ratio 1:2) peptides. Gray columns indicate elution volume of the peptides alone from panel B.

##### Fig S5. ON/OFF AAV-targeting strategy in Hoxb8-Cre mice.

**A)** Layout showing peptide expression in WT mice of mixed genders using an ON-OFF viral targeting approach. With this, peptide expression can be restricted to neurons expressing Cre-recombinase or not, as indicated.

**B)** Same as for panel A, but for Hoxb8-Cre mice.

**C)** Paw withdrawal threshold (von Frey) before and after i.t. injection of AAVs and after intraplantar CFA administration in Hoxb8-Cre and WT mice using the ON-OFF strategy. $^{**}P < 0.01$, $^{****}P < 0.0001$. Two-way repeated measures ANOVA, $F(9, 93) = 3.800$; Tukey's multiple comparisons; $N$ = 7-10 per group.

**D)** Day 4 from experiment in panel C.

Abbreviations: BL, baseline; i.t., intrathecal. Data in C and D are shown as mean with s.e.m.

##### Fig S6. Robust AAV transduction of dorsal column fibers following intrathecal injection of AAV8 into Ai14 mice.

**A)** Ai14 transgenic mice harboring a lox-stop-lox (LSL)-tdTomato cassette controlled by the CAG promoter in the Rosa26 locus (Rosa-CAG-LSL-tdTomato) were i.t. injected with AAV8-hSyn-iCre:EGFP-P2A-Di-C5. The Cre-dependent tdTomato fluorescent signal allows for precise identification of cells and circuits transduced by the injected AAV vector.

**B)** Whole-mount imaging showing part of the thoracic spinal cord region labeled by tdTomato with noticeable fibers along the spinal cord dorsal column. $N$ = 8.

**C)** tdTomato-positive fibers are visible within the spinal cord dorsal column at the sacral, lumbar, thoracic, and cervical levels. These fibers originate from DRG neurons. The AAV has limited access to the brain with sparse tdTomato-positive neurons found across different brain regions. This sagittal brain slice example showing limited transduction of neurons within the cerebellum and olfactory system. This sparse labeling profile of the brain was consistent but varied across regions in different animals. *N* = 9. All scalebars represent 500 µm.

**D)** Paw withdrawal threshold (von Frey) measured before and after intraplantar CFA administration in Ai14 reporter mice i.t. injected with AAV8-iCre:EGFP-P2A-Di-C5. Two-way repeated measures ANOVA, F(2, 12) = 3.146; Holm-Šidák's multiple comparisons; *N* = 4 per group, mixed genders. Data are shown as mean with s.e.m. Abbreviations: BL, baseline.

**E)** Representative images of neuronal cell bodies in the DRG show immunostaining with markers IB4, CGRP, and NF200 (top panels) with tdTomato as a marker for viral transduction (middle panels) and merged images (bottom panels). Arrows mark double-positive neurons.

**F)** Histogram displaying numbers and cross-sectional area of neuronal cell bodies from L3 and L4 DRG from Panel E. All neurons = 1367, tdTomato-positive neurons = 224. Insert: Cumulative distribution of all neurons and tdTomato-positive neurons as a function of their cross-sectional area. *P* < 0.0001; Kolmogorov-Smirnov test.

**G)** The relative distribution of all neurons (solid grey area/line) from Panel E plotted together with the relative distribution of all transduced neurons (solid red area) as a function of their cross-sectional area. The dotted lines show the cross-sectional area of neurons positive for IB4, CGRP, and NF200 markers, respectively. *N* = 40-44 neurons per marker, *N* = 3 mice for panel E-G.

#### Fig. S7. Confirmation of treatment in SNI animals and assessment of general behavior and locomotion.

**A)** Study layout and timeline.

**B)** Mechanical paw withdrawal threshold (von Frey) in WT male mice before and after SNI or sham surgery with i.t. AAV injection with AAV8-Di(7P14P)-C5 or AAV8-tdTomato performed 2 days post SNI or sham surgery. Two-way ANOVA, F(6, 38) = 9.390; Tukey's multiple comparisons; *N* = 5-6 per group. Abbreviations: BL, baseline; i.t., intrathecal.

**C-I)** General behaviors were monitored for 22 hours on days 55-58 after SNI or sham surgery using individual Laboras-specific cages and monitoring system. All comparisons were one-way ANOVA with Tukey's multiple comparisons and were all non-significant. *N* = 5-6 per group.

**J)** Max contact area of the ipsilateral paw determined by CatWalk in WT mice before and after SNI or sham surgery with i.t. AAV injections of AAV8-Di(7P14P)-C5 or AAV8-tdTomato performed 2 days post SNI or sham surgery. Mean of three trials at each time point. Two-way repeated measures ANOVA, F(9, 57) = 6.855; Tukey's multiple comparisons; *N* = 5-6 per group.

**K)** As in panel J, but for the swing speed of the ipsilateral paw. Mean of three trials at each time point. Two-way repeated measures ANOVA, F(9, 57) = 2.140; Tukey's multiple comparisons; *N* = 5-6 per group.

**L)** As in panel J, but for average run speed. Mean of three trials at each time point. Two-way repeated measures ANOVA, F(9, 57) = 0.9906; Tukey's multiple comparisons; *N* = 5-6 per group.

**M)** As in panel J, but for the run duration. Mean of three trails at each time point. Two-way repeated measures ANOVA, F(9, 57) = 0.7267; Tukey's multiple comparisons; *N* = 5-6 per group.

**N)** Average distance of voluntary wheel running during the night-time (18:00-06:00) as measured 3 times in the same animals (SNI and sham-operated mice with AAV8-Di(7P14P)-C5 or AAV8-tdTomato). One-way ANOVA, F(9, 19) = 10.32; Tukey's multiple comparison; *N* = 5-6 per group.

**O)** Body weight measured on multiple occasions from the day of SNI surgery in SNI and sham-operated mice with AAV8-Di(7P14P)-C5 or AAV8-tdTomato. $^{**}P < 0.01$ (comparing sham w. AAV8-Di(7P14P)-C5 vs. SNI w. AAV8-tdTomato from day 33-91). Two-way repeated measures ANOVA, F(30, 190) = 2.703; Tukey's multiple comparisons; *N* = 5-6 per group.

Data in B-O are shown as mean with s.e.m. Abbreviations: BL, baseline; ipsi, ipsilateral.

**Fig. S8. Time course of open field test and validation of AAV treatments.**

**A)** Study layout and timeline.

**B)** Two hours open field test (OFT) showing distance traveled (5 min interval binned) for AAV8-Di-C5, AAV8-tdTomato, and vehicle-treated mice. Two-way repeated measures ANOVA, $F(46, 299) = 1.097$; Dunnett's multiple comparisons; $N$ = 5-6 per group.

**C)** Validation of the treatments in the CFA model following the behavioral assays. Two-way ANOVA, $F(2, 13) = 7.428$; Tukey's multiple comparisons; $N$ = 5-6 per group.

Abbreviations: i.t., intrathecal. All data in B-C are shown as mean with s.e.m.

#### Table S1. Quantitative real-time PCR (qRT-PCR) of mRNA transcription of peptide and housekeeping genes in various tissues.

Mean Ct values with standard deviation ($N$ = 4 animals, 3 technical replicates each) are shown for each primer target and tissue type. Mean Ct values are subsequently normalized to the expression of housekeeping genes (Methods, qPCR, equation 1) (values not shown). The relative change in peptide expression compared to the control group, DPBS, is expressed as mean ΔΔCt values with standard deviation for each treatment group (Methods, qPCR, equation 2). P-values are shown (unpaired t-test) when comparing the relative expression (expressed as ΔΔCt) in the treatment group, AAV8-Di-C5, to the control group, DPBS.

### Suppl. Fig 1

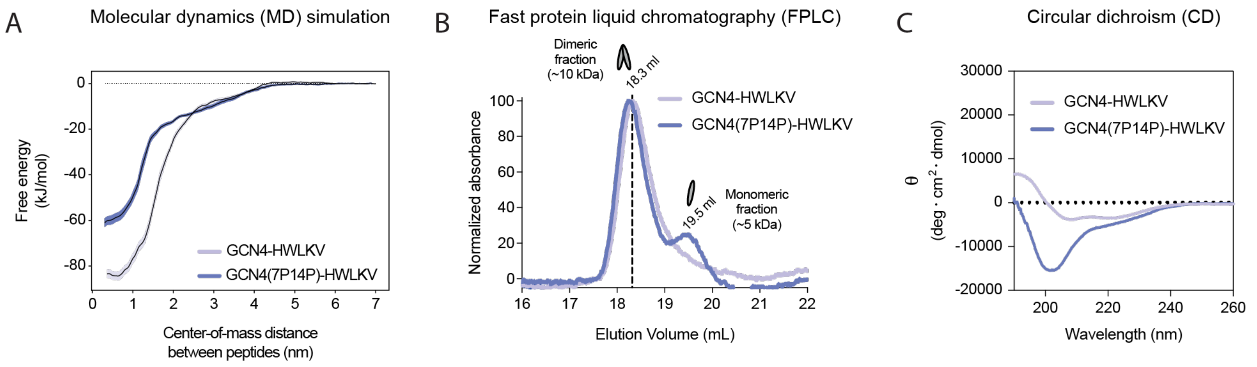


### Suppl. Fig 2

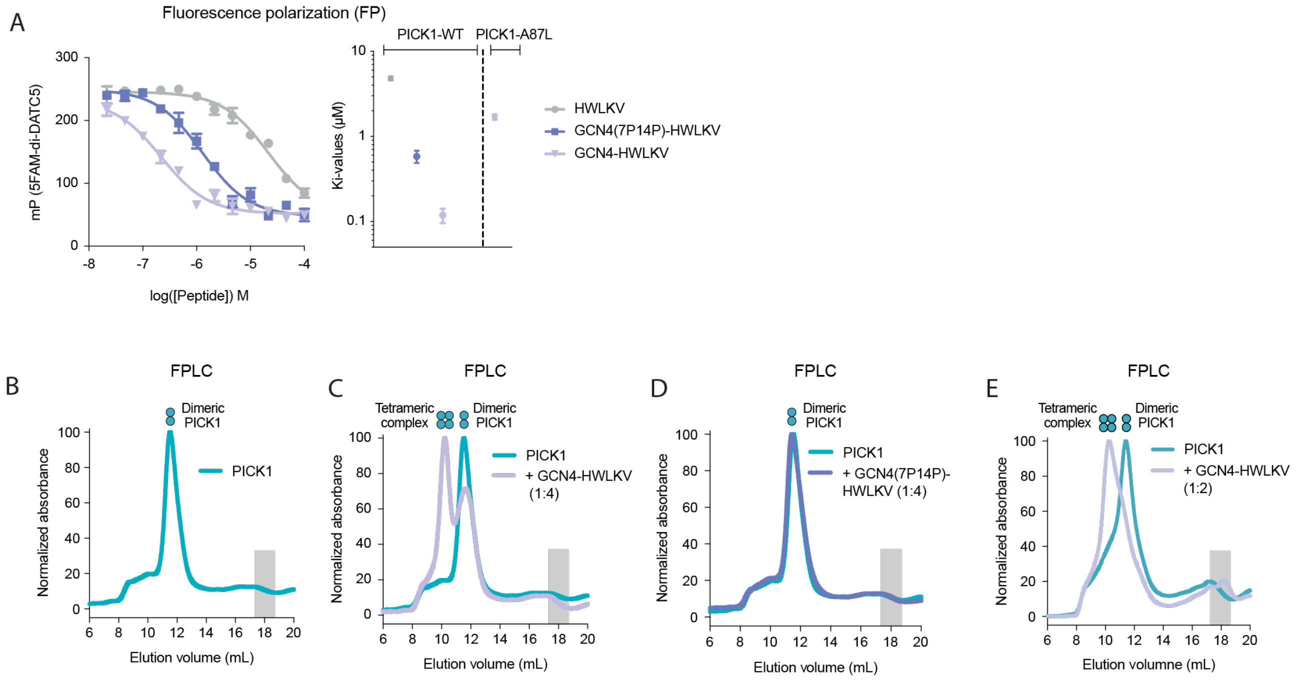

### Suppl. Fig 3

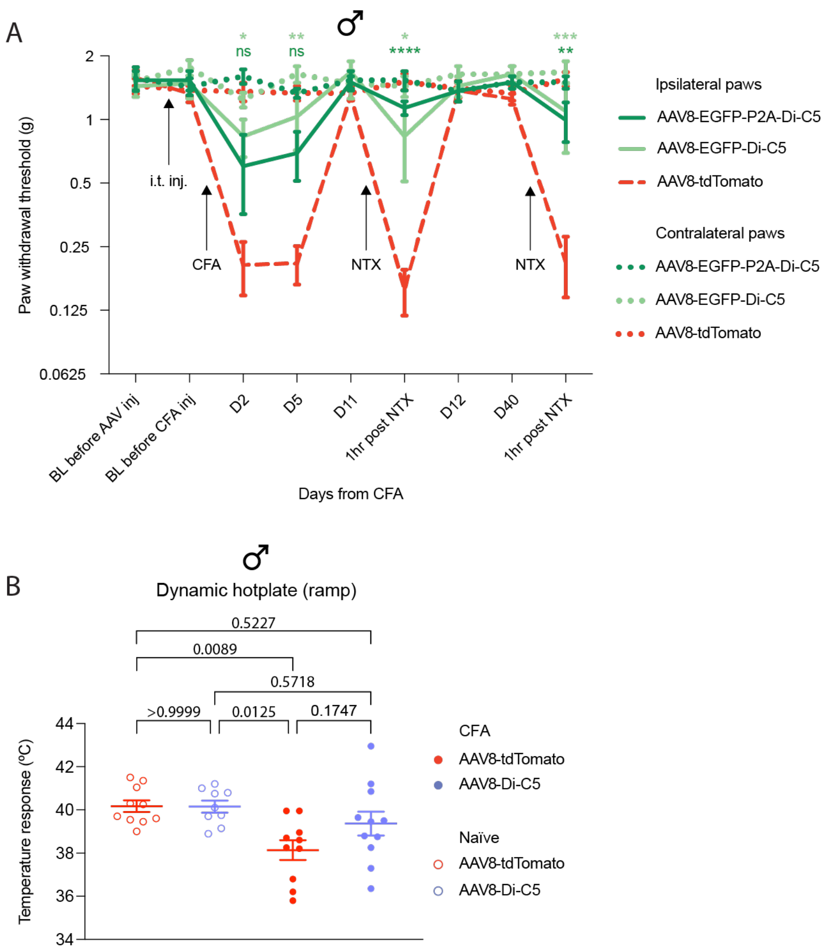
A
♂
Paw withdrawal threshold (g)
2
1
0.5
0.25
0.125
0.0625
ns
ns
****
**
i.t. inj.
CFA
NTX
NTX
BL before AAV inj
BL before CFA inj
D2
D5
D11
1hr post NTX
D12
D40
1hr post NTX
Days from CFA
Ipsilateral paws
AAV8-EGFP-P2A-Di-C5
AAV8-EGFP-Di-C5
AAV8-tdTomato
Contralateral paws
AAV8-EGFP-P2A-Di-C5
AAV8-EGFP-Di-C5
AAV8-tdTomato
B
♂
Dynamic hotplate (ramp)
0.5227
0.0089
0.5718
>0.9999
0.0125
0.1747
Temperature response (ºC)
44
42
40
38
36
34
CFA
AAV8-tdTomato
AAV8-Di-C5
Naïve
AAV8-tdTomato
AAV8-Di-C5

### Suppl. Fig 4

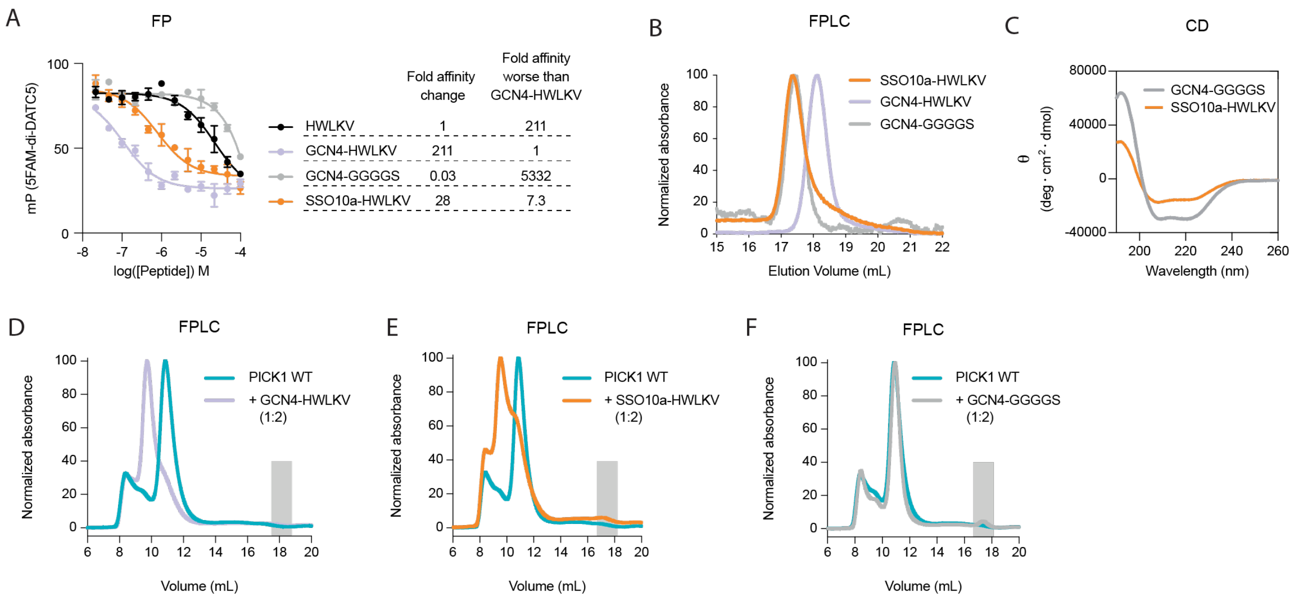
A
FP
mP (5FAM-di-DATC5)
log([Peptide]) M
Fold affinity change
Fold affinity worse than GCN4-HWLKV
HWLKV 1 211
GCN4-HWLKV 211 1
GCN4-GGGGS 0.03 5332
SSO10a-HWLKV 28 7.3
B
FPLC
Normalized absorbance
Elution Volume (mL)
SSO10a-HWLKV
GCN4-HWLKV
GCN4-GGGGS
C
CD
θ (deg · cm² · dmol)
Wavelength (nm)
GCN4-GGGGS
SSO10a-HWLKV
D
FPLC
Normalized absorbance
Volume (mL)
PICK1 WT
+ GCN4-HWLKV (1:2)
E
FPLC
Normalized absorbance
Volume (mL)
PICK1 WT
+ SSO10a-HWLKV (1:2)
F
FPLC
Normalized absorbance
Volume (mL)
PICK1 WT
+ GCN4-GGGGS (1:2)

### Suppl. Fig 5

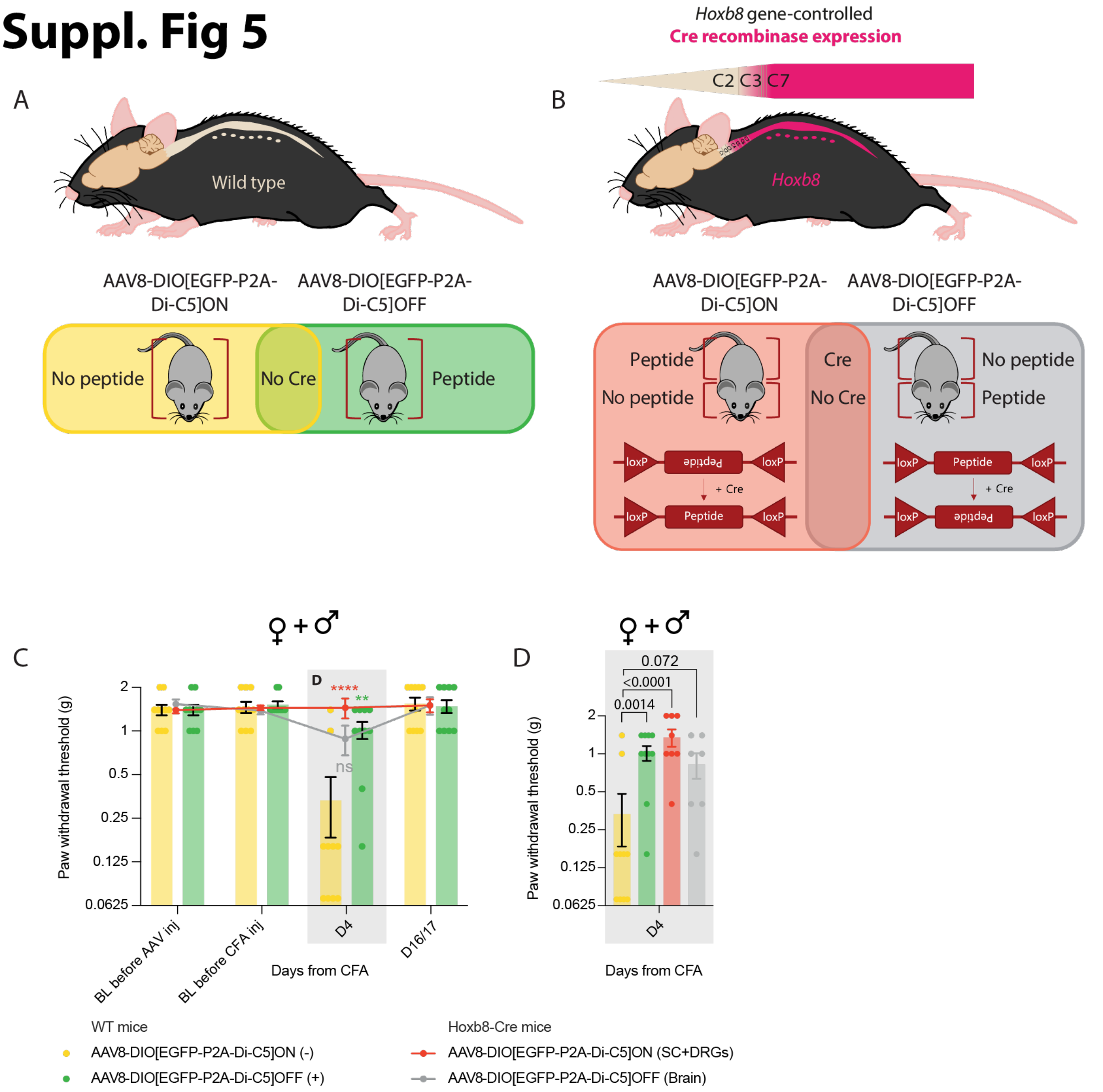
A
Wild type
AAV8-DIO[EGFP-P2A-Di-C5]ON
AAV8-DIO[EGFP-P2A-Di-C5]OFF
No peptide
No Cre
Peptide
B
Hoxb8 gene-controlled
Cre recombinase expression
C2 C3 C7
Hoxb8
AAV8-DIO[EGFP-P2A-Di-C5]ON
AAV8-DIO[EGFP-P2A-Di-C5]OFF
Peptide
No peptide
Cre
No Cre
No peptide
Peptide
loxP
Peptide
+ Cre
C
♀+♂
Paw withdrawal threshold (g)
2
1
0.5
0.25
0.125
0.0625
D
****
**
ns
BL before AAV inj
BL before CFA inj
D4
D16/17
Days from CFA
D
♀+♂
0.072
<0.0001
0.0014
Paw withdrawal threshold (g)
D4
Days from CFA
WT mice
AAV8-DIO[EGFP-P2A-Di-C5]ON (-)
AAV8-DIO[EGFP-P2A-Di-C5]OFF (+)
Hoxb8-Cre mice
AAV8-DIO[EGFP-P2A-Di-C5]ON (SC+DRGs)
AAV8-DIO[EGFP-P2A-Di-C5]OFF (Brain)

### Suppl. Fig 6

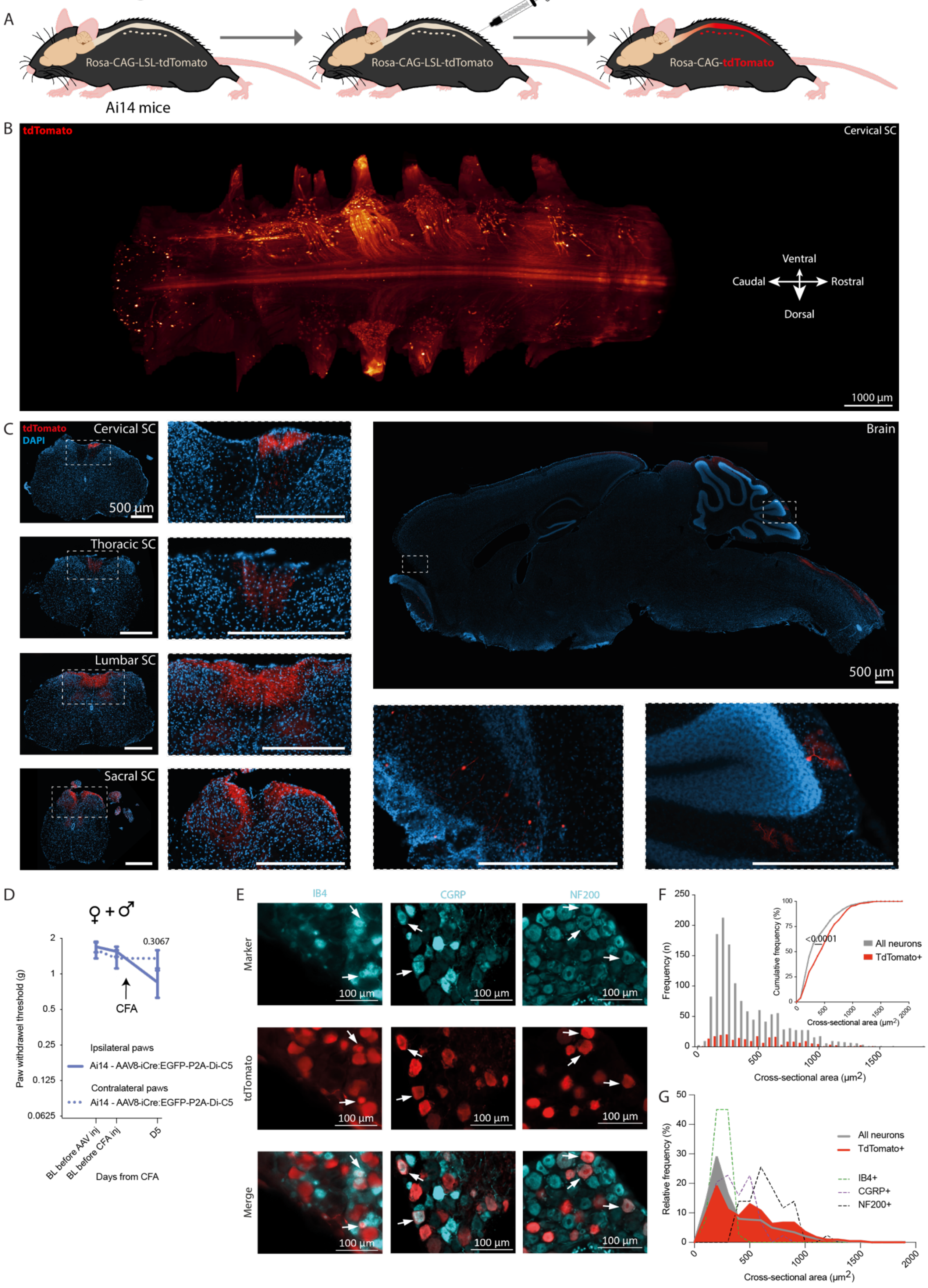

A
AAV2.8-hSyn-iCre:EGFP-P2A-Di-C5
Rosa-CAG-LSL-tdTomato
Rosa-CAG-LSL-tdTomato
Rosa-CAG-tdTomato
Ai14 mice
B
tdTomato
Cervical SC
Ventral
Caudal
Rostral
Dorsal
1000 µm
C
tdTomato
DAPI
Cervical SC
500 µm
Thoracic SC
Lumbar SC
Sacral SC
Brain
500 µm
D
♀+♂
Paw withdrawel threshold (g)
0.3067
CFA
Ipsilateral paws
Ai14 - AAV8-iCre:EGFP-P2A-Di-C5
Contralateral paws
Ai14 - AAV8-iCre:EGFP-P2A-Di-C5
BL before AAV inj
BL before CFA inj
D5
Days from CFA
E
IB4
CGRP
NF200
Marker
tdTomato
Merge
100 µm
F
Frequency (n)
Cumulative frequency (%)
<0.0001
All neurons
TdTomato+
Cross-sectional area (µm2)
Cross-sectional area (µm2)
G
Relative frequency (%)
All neurons
TdTomato+
IB4+
CGRP+
NF200+
Cross-sectional area (µm2)

### Suppl. Fig 7

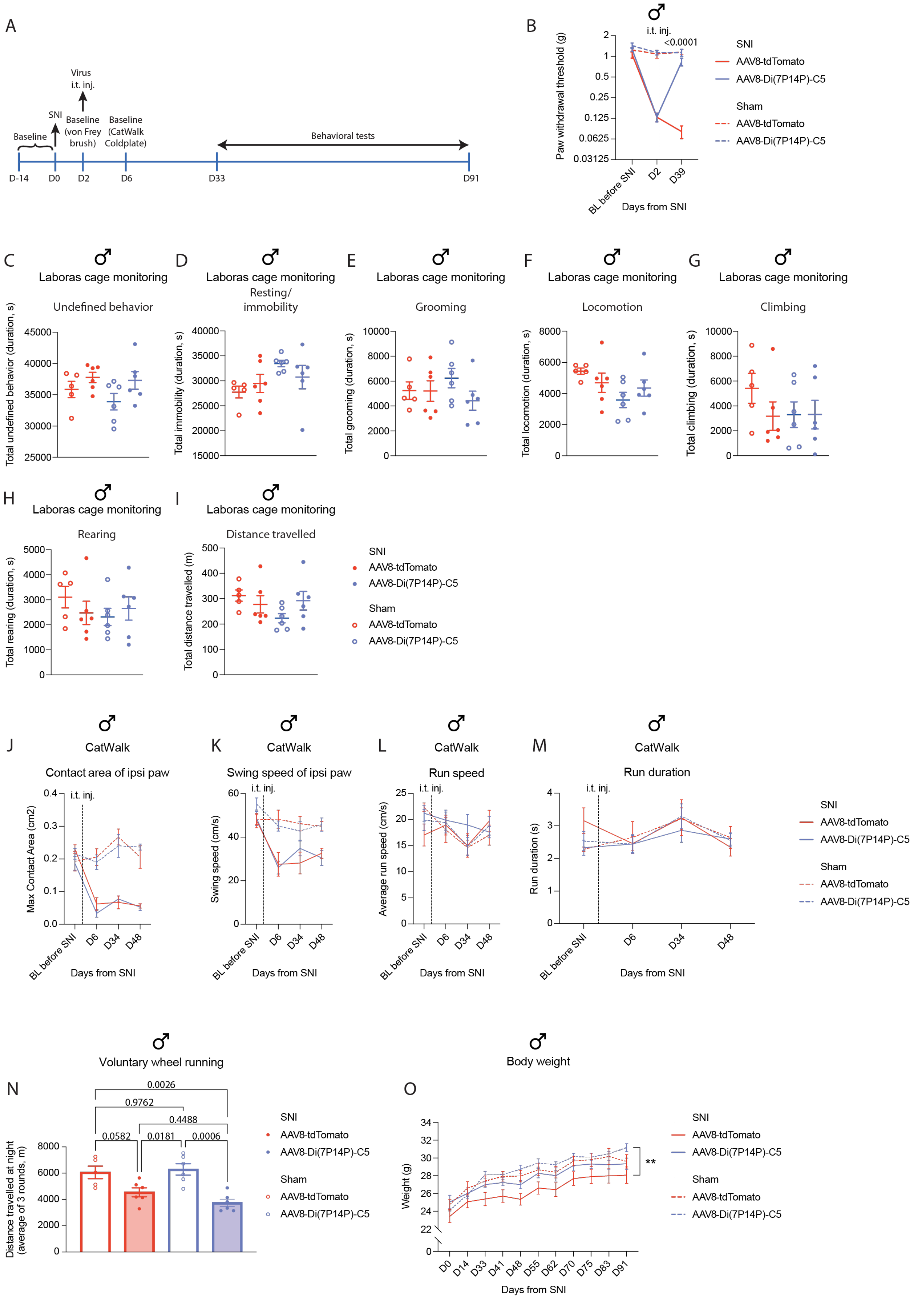
A
Virus
i.t. inj.
SNI
Baseline
Baseline
(von Frey
brush)
Baseline
(CatWalk
Coldplate)
Behavioral tests
D-14
D0
D2
D6
D33
D91
B
i.t. inj.
<0.0001
Paw withdrawal threshold (g)
BL before SNI
D2
D39
Days from SNI
SNI
AAV8-tdTomato
AAV8-Di(7P14P)-C5
Sham
AAV8-tdTomato
AAV8-Di(7P14P)-C5
C
Laboras cage monitoring
Undefined behavior
Total undefined behavior (duration, s)
D
Laboras cage monitoring
Resting/
immobility
Total immobility (duration, s)
E
Laboras cage monitoring
Grooming
Total grooming (duration, s)
F
Laboras cage monitoring
Locomotion
Total locomotion (duration, s)
G
Laboras cage monitoring
Climbing
Total climbing (duration, s)
H
Laboras cage monitoring
Rearing
Total rearing (duration, s)
I
Laboras cage monitoring
Distance travelled
Total distance travelled (m)
SNI
AAV8-tdTomato
AAV8-Di(7P14P)-C5
Sham
AAV8-tdTomato
AAV8-Di(7P14P)-C5
J
CatWalk
Contact area of ipsi paw
i.t. inj.
Max Contact Area (cm2)
Days from SNI
K
CatWalk
Swing speed of ipsi paw
i.t. inj.
Swing speed (cm/s)
Days from SNI
L
CatWalk
Run speed
i.t. inj.
Average run speed (cm/s)
Days from SNI
M
CatWalk
Run duration
i.t. inj.
Run duration (s)
Days from SNI
BL before SNI
D6
D34
D48
SNI
AAV8-tdTomato
AAV8-Di(7P14P)-C5
Sham
AAV8-tdTomato
AAV8-Di(7P14P)-C5
N
Voluntary wheel running
0.0026
0.9762
0.4488
0.0582
0.0181
0.0006
Distance travelled at night
(average of 3 rounds, m)
SNI
AAV8-tdTomato
AAV8-Di(7P14P)-C5
Sham
AAV8-tdTomato
AAV8-Di(7P14P)-C5
O
Body weight
Weight (g)
**
D0
D14
D33
D41
D48
D55
D62
D70
D75
D83
D91
Days from SNI
SNI
AAV8-tdTomato
AAV8-Di(7P14P)-C5
Sham
AAV8-tdTomato
AAV8-Di(7P14P)-C5

### Suppl. Fig 8

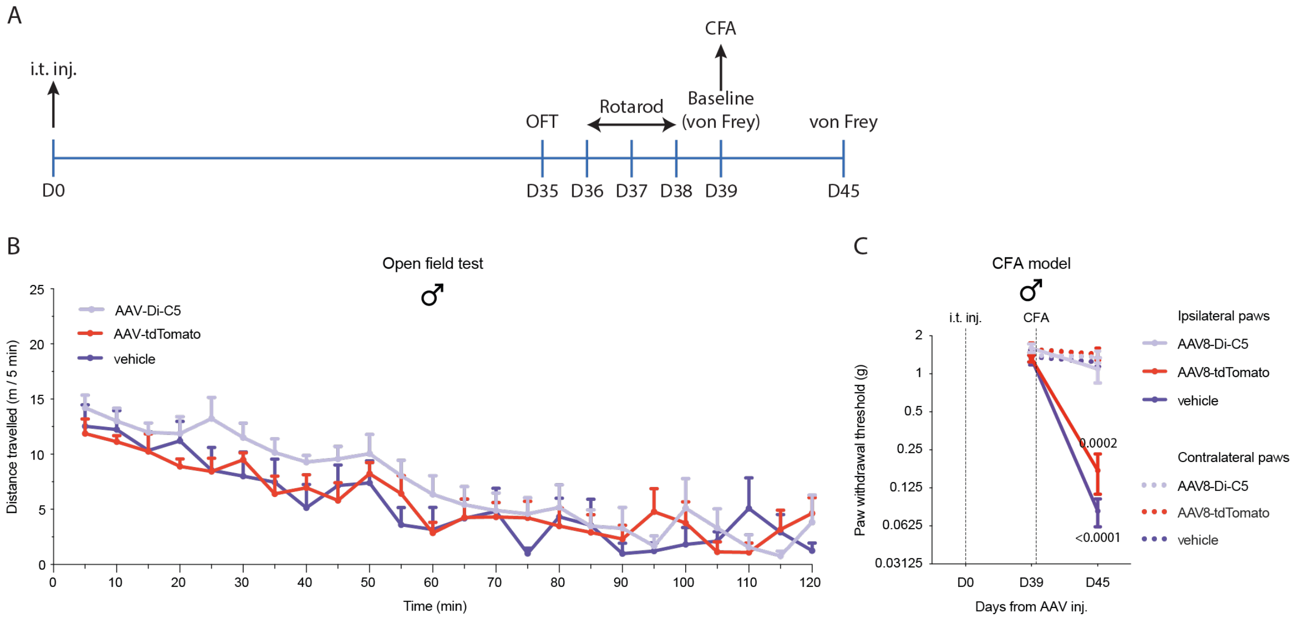
A
i.t. inj.
CFA
Rotarod
Baseline
(von Frey)
OFT
von Frey
D0
D35
D36
D37
D38
D39
D45
B
Open field test
AAV-Di-C5
AAV-tdTomato
vehicle
Distance travelled (m / 5 min)
Time (min)
C
CFA model
i.t. inj.
CFA
Paw withdrawal threshold (g)
0.0002
<0.0001
Days from AAV inj.
D0
D39
D45
Ipsilateral paws
AAV8-Di-C5
AAV8-tdTomato
vehicle
Contralateral paws
AAV8-Di-C5
AAV8-tdTomato
vehicle
