## Supplementary Table 1 for "Recombinant dimeric PDZ protein inhibitors for long-term relief of chronic pain by AAV therapeutics"

| Tissue | Primer target | AAV-Di-C5<br>mean Ct ± SD[¥] | DPBS<br>mean Ct ± SD[¥] | mean ΔΔCt ± SD (AAV-Di-C5) | mean ΔΔCt ± SD (DPBS) | p-value (Unpaired t-test) |
|---|---|---|---|---|---|---|
| Dorsal root ganglion | Di-C5 | 24.38 ± 1.22 | 34,63 ± 0,43 | 10,15 ± 1,32 | 0,0025 ± 0,78 | <0,0001 |
| | *EEF1A1* | 18,18 ± 0,31 | 18,39 ± 0,66 | | | |
| | *YHWAS* | 18,41 ± 0,29 | 18,56 ± 0,66 | | | |
| Sciatic nerve | Di-C5 | 33,19 ± 1,33 | 34,66 ± 0,40 | 1,62 ± 0,81 | 0,0025 ± 0,97 | 0,04 |
| | *EEF1A1* | 19,87 ± 0,88 | 19,71 ± 0,87 | | | |
| | *YHWAS*[†] | - | - | | | |
| Dorsal column | Di-C5 | 33,16 ± 1,86 | 33,44 ± 3,11 | 0,03 ± 1,63 | 0,000 ± 2,92 | 0,99 |
| | *EEF1A1* | 18,12 ± 0,20 | 18,36 ± 0,56 | | | |
| | *YHWAS* | 18,95 ± 0,31 | 19,21 ± 0,44 | | | |
| Lumbar spinal cord | Di-C5 | 32,52 ± 1,48 | 34,68 ± 0,64 | 2,08 ± 1,22 | -0,0025 ± 0,58 | 0,02 |
| | *EEF1A1* | 18,30 ± 0,33 | 18,43 ± 0,32 | | | |
| | *YHWAS* | 19,30 ± 0,36 | 19,34 ± 0,31 | | | |
| Cervical spinal cord | Di-C5 | 33,73 ± 1,43 | 34,84 ± 0,32 | 1,11 ± 1,20 | 0,000 ± 0,30 | 0,12 |
| | *EEF1A1* | 18,50 ± 0,40 | 18,53 ± 0,60 | | | |
| | *YHWAS* | 19,39 ± 0,44 | 19,36 ± ,45 | | | |
| Pons | Di-C5 | 34,32 ± 0,94 | 34,17 ± 1,20 | -0,30 ± 0,97 | -0,0025 ± 1,63 | 0,77 |
| | *EEF1A1* | 18,74 ± 0,59 | 19,22 ± 0,76 | | | |
| | *YHWAS* | 19,63 ± 0,51 | 20,01 ± 0,70 | | | |
| Cerebellum | Di-C5 | 34,05 ± 0,78 | 34,35 ± 0,88 | 0,85 ± 0,58 | 0,0025 ± 0,93 | 0,17 |
| | *EEF1A1* | 18,47 ± 0,84 | 17,90 ± 0,55 | | | |
| | *YHWAS* | 19,37 ± 0,76 | 18,84 ± 0,66 | | | |
| Liver | Di-C5 | 25,91 ± 0,78 | 35,00 ± 0,00 | 9,43 ± 0,20 | 0,000 ± 0,39 | <0,0001 |
| | *EEF1A1* | 16,62 ± 0,67 | 16,48 ± 0,50 | | | |
| | *YHWAS* | 21,25 ± 0,67 | 20,72 ± 0,30 | | | |

[¥] each mean Ct value is a mean of 3 technical replicates from each of 4 different animals with a threshold of 35

[†] for the sciatic nerve, only the housekeeping gene *EEF1A1* was used for normalization, as the housekeeping gene *YHWAS* exhibited high Ct variation
